# Oncometabolite-driven enhancer hypermethylation promotes a hyporesponsive microglial state in IDH-mutant gliomas

**DOI:** 10.1101/2024.08.23.608811

**Authors:** Alice Laurenge, Pietro Pugliese, Sarah Scuderi, Bertrand Mathon, Louisa Drouiche, Quentin Richard, Stéphanie Jouannet, Chaima Benmohamed, Kirill Smirnov, Lea Hanna Doumit Sakr, Yvette Hayat, Nina Pottier, Pauline Marijon, Karim Labreche, Agustí Alentorn, Maïté Verreault, Ahmed Idbaih, Cristina Birzu, Aurore Desmons, Inès Fayache, Philip J. Kingsley, Lawrence J. Marnett, Karima Mokhtari, Suzanne Tran, Emmanuelle Huillard, Eric Duplus, Elias El-Habr, Jaydutt Bhalshankar, Lucas A. Salas, Mario L. Suvà, Michele Ceccarelli, Antonio Iavarone, Gilles Huberfeld, Garrett A. Kaas, Franck Bielle, Mehdi Touat, Michel Mallat, Marc Sanson, Luis Jaime Castro-Vega

## Abstract

Tumor-associated microglia and macrophages (TAMs), the most abundant myeloid populations in gliomas, shape immune responses through transcriptional programs influenced by the tumor microenvironment. Although these programs differ according to tumor IDH status, the underlying epigenetic mechanisms remain poorly understood. Here, we uncover widespread DNA hypermethylation in the myeloid compartment of IDH-mutant gliomas, predominantly at distal enhancers enriched for motifs of core microglial transcription factors (TFs). This remodeled enhancer landscape strongly correlated with reduced activity of TF regulons and coordinated repression of immunomodulatory programs that normally support microglial activation. Using primary human microglia, we show that prolonged exposure to the oncometabolite D-2-hydroxyglutarate (D-2HG) reduces TET activity and increases 5mC/5hmC ratios near TF-binding motifs within enhancers affected *ex vivo.* Consistent with these epigenetic alterations, D-2HG-treated microglia exhibited transcriptional signatures compatible with blunted proinflammatory responses, whereas pharmacological inhibition of mutant IDH in patients partially restored microglial immune reactivity. Altogether, our findings reveal a chronic D-2HG–driven epigenetic priming mechanism that promotes a hyporesponsive microglial state, providing a rationale for the immunologically cold phenotype of IDH-mutant gliomas and offering insight into how IDH-targeted therapies may reshape microglial immune responses.

## Introduction

Diffuse gliomas are the most common primary brain tumors that invariably recur and remain incurable despite current standard therapies and immune checkpoint inhibition^1,2^. A large proportion of these tumors harbor mutations in the isocitrate dehydrogenase 1 (*IDH1*) or 2 (*IDH2*), early initiating events in gliomagenesis, which confer a neomorphic enzymatic ability converting α-ketoglutarate (α-KG) into D-2-hydroxyglutarate (D-2HG)^3^. D-2HG promotes cellular transformation by inhibiting α-KG-dependent dioxygenases, including the TET family of DNA demethylases and KDM histone demethylases, thereby driving widespread epigenomic dysregulation in tumor cells^4^.

Although the cell-intrinsic effects of D-2HG have been extensively characterized, its oncogenic influence on the tumor microenvironment (TME), where this oncometabolite also accumulates^5,6^, remains poorly understood. Emerging evidence shows that D-2HG can suppress T-cell activation and alter the metabolism of macrophages, dendritic cells, and CD8^+^ T cells, mostly through non-epigenetic mechanisms^7–13^. The impact of D-2HG on microglia, the main innate immune cells of the brain that dominate the myeloid compartment of IDH-mutant gliomas and orchestrate inflammatory responses, remains to be determined. As long-lived tissue-resident cells, microglia are chronically exposed to D-2HG throughout glioma development.

Tumor-associated macrophages and microglia (TAMs) constitute up to 40% of the tumor mass and play critical roles in glioma progression^14^. These cells are therefore recognized as prognostic biomarkers and potential therapeutic targets^15,16^. TAMs are highly plastic, and their phenotypes and immune responses are shaped by both ontogeny and microenvironmental cues^17–19^. In IDH-wt glioma models, resident microglia and monocyte-derived macrophages (MDMs) display distinct transcriptional and chromatin landscapes^20^. Similarly, transcriptomic differences have been observed in myeloid cells from human gliomas as a function of the IDH status^21,22^. In IDH-wt tumors, both MDMs and microglia exhibit activation-associated expression profiles, including elevated expression of MHC class genes, whereas in IDH-mutant gliomas they more closely resemble homeostatic microglia from non-tumor brain tissue^21^. Single-cell RNA-seq studies have further delineated subsets of MDMs and microglia, some correlating with clinical outcome^23–30^.

Most recently, it was shown that myeloid cells in human gliomas express shared immunomodulatory activity programs that transcend classical lineage boundaries^31^. These programs are regulated by stimulus-induced transcription factors, with AP-1 increasingly dominating over NF-κB as tumor progress, driving a shift from inflammatory to immunosuppressive states^31^. However, the epigenetic mechanisms that define or stabilize these programs remain largely uncharacterized. Moreover, while hypoxia is a well-established driver of the immunosuppressive TAM phenotype in IDH-wt tumors^30–32^, little is known about the signals modulating the functional states of TAMs in the IDH-mutant microenvironment.

Given the high concentrations of D-2HG in IDH-mutant gliomas, we hypothesized that chronic oncometabolite signaling in the TME may alter DNA methylation at cis-regulatory elements in tissue-resident microglia, thereby affecting immune-related gene expression programs. To test this, we integrated DNA methylome, transcriptome, and targeted proteomic analyses of CD11B⁺ myeloid cells isolated from human gliomas and non-tumor brain tissues, complemented by epigenomic and functional analyses in primary human microglia and an *in vivo* D-2HG delivery model.

Our findings reveal that CD11B⁺ myeloid cells in IDH-mutant gliomas exhibit pervasive DNA hypermethylation at enhancer elements associated with reduced activity of lineage-determining and stimulus-responsive TFs that govern microglial activation. Consequently, these cells display a broadly hyporesponsive state, marked by attenuation of key activation-associated gene networks. Of note, primary human microglia and murine CD11B⁺ myeloid cells exposed to D-2HG *in vivo* exhibit methylation changes compatible with impaired methylation turnover at enhancers identified as hypermethylated in the myeloid compartment of IDH-mutant gliomas. Consistent with these epigenetic changes, D-2HG-treated microglia repress activation-associated gene programs and display blunted inflammatory responses following LPS stimulation, thereby recapitulating key features of the *ex vivo* phenotype. Importantly, treatment with mutant IDH inhibitors in patients partially restored microglial immune-related transcriptional programs linked to enhancers hypermethylated in tumors. Together, these findings support a model in which D-2HG-driven epigenetic reprogramming contributes to suppression of resident microglia as an early mechanism of immune evasion in IDH-mutant gliomas, providing a framework to understand their immunologically cold TME and how it can be modulated by IDH-targeted therapies.

## Results

### Study design

To investigate the epigenetic basis of TAM phenotypes associated with IDH status, we profiled the bulk DNA methylome and/or transcriptome of CD11B⁺ myeloid cells isolated by magnetic sorting from adult-type diffuse gliomas (*n* = 54), including IDH-wt glioblastomas, IDH-mutant oligodendrogliomas, and IDH-mutant astrocytomas **(Fig. 1A)**. We also analyzed publicly available datasets of CD11B⁺ cells from non-tumor brain tissues (*n* = 14), isolated using similar procedures and profiled with the same technologies^33,34^. Targeted proteomics and LC-MS/MS assays of global DNA methylation were performed in independent cohorts of tumors and non-tumor tissues (*n* = 29) to complement these datasets. The pathological, molecular and demographic characteristics of the study cohort are detailed in **Supplementary Table S1**. Contamination of CD11B⁺ fractions by tumor cells was evaluated by detecting *IDH1* R132H or TERT promoter mutations (C228T/C250T) using droplet digital PCR (ddPCR) and further assessed through RNA-seq-based estimation of *IDH* variant allelic frequency **(Supplementary Fig. S1A and S1B)**. Together, these approaches indicated minimal tumor-cell contamination in the CD11B⁺ fractions used for downstream studies, with estimated purity reaching ∼95%. These analyses also ruled out the possibility that TAMs from our study cohort harbored the IDH1 R132H mutation to a significant extent, as previously reported^35^.

**Figure 1.**
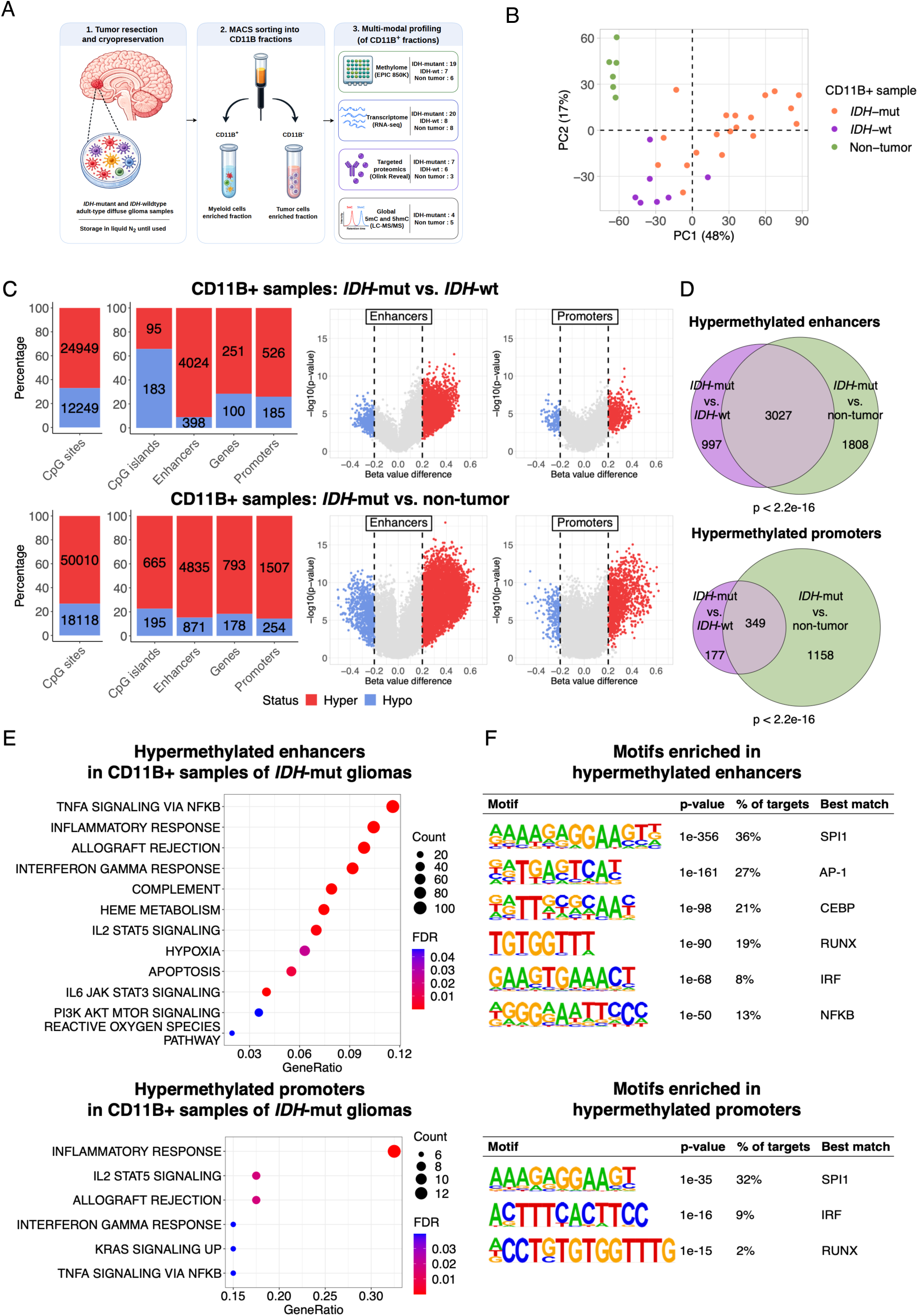
Widespread DNA hypermethylation in CD11B^+^ myeloid cells from IDH-mutant gliomas. **(A)** Overview of the *ex vivo* study workflow. Glioma samples underwent enzymatic dissociation followed by CD11B-based magnetic sorting to isolate myeloid cell-enriched and tumor cell-enriched fractions. DNA and/or RNA extracted from CD11B⁺ fractions were used for downstream methylome and transcriptome analyses, whereas available datasets from non-tumor brain tissues were included as controls for comparative analyses. Orthogonal validation experiments performed in independent CD11B⁺ fractions from tumors and non-tumor tissues included targeted proteomics and global 5mC and 5hmC quantification. **(B)** Principal component analysis performed using methylation profiles from approximately 600,000 CpG sites across the genome in CD11B^+^ fractions from the study cohort. **(C)** Stacked bar plots (left) showing the absolute number and relative proportion of differentially methylated CpG sites across four genomic regions according to methylation status (absolute Δβ > 0.2, FDR < 0.05) for the indicated comparisons. Volcano plots (right), colored according to methylation status, illustrate the magnitude and distribution of differentially methylated enhancers and promoters in the indicated comparisons. **(D)** Venn diagrams showing significant overlaps between hypermethylated enhancers (upper panel) or promoters (lower panel) identified in CD11B^+^ fractions from IDH-mutant gliomas relative to either IDH-wt tumors or non-tumor brain tissues. P-values were calculated using over-representation analysis (ORA). **(E)** Bubble plots showing Hallmark gene sets enriched among genes associated with hypermethylated enhancers or promoters across comparisons in CD11B^+^ fractions from IDH-mutant gliomas. Dot size and color indicate the number of genes contributing to each category and the corresponding FDR value, respectively (FDR < 0.05). **(F)** Significant DNA sequence motifs (ranked by p-value) enriched within hypermethylated enhancers or promoters across comparisons in CD11B^+^ fractions from IDH-mutant gliomas were identified using HOMER.

### CD11B⁺ myeloid cells in IDH-mutant gliomas display global DNA hypermethylation

To establish the DNA methylation landscape of our study cohort in relation to IDH status, we first profiled tumor cells enriched fractions (CD11B⁻) isolated from both IDH-wt and IDH-mutant gliomas (*n* = 17, Supplementary Table S1) using MethylationEPIC (EPIC) arrays. Genome-wide analysis of CpG methylation confirmed that IDH-mutant gliomas exhibit global hypermethylation, albeit with considerable intertumoral heterogeneity, consistent with previous reports^36–38^ **(Supplementary Fig. S2A and S2B)**. In total, 49,676 CpG sites were differentially methylated (absolute Δβ > 0.2, FDR < 0.05), with 92.5% of these hypermethylated in IDH-mutant compared to IDH-wt tumors **(Supplementary Fig. S2C)**. Mapping of these CpGs across functional genomic regions revealed enrichment in promoters and enhancers. Gene promoters were defined as sequences within −1.5 kb to +0.5 kb of the transcription start sites (TSS), whereas enhancers were defined as distal regulatory elements linked to promoter regions within 500 kb (FDR < 1 x 10^−5^; Pearson’s correlation), following the FANTOM5 consortium framework^39^. This analysis confirmed that enhancers are particularly sensitive to hypermethylation in IDH-mutant glioma cells^40^ **(Supplementary Fig. S2C and S2D)**. We next examined methylome changes in CD11B⁺ myeloid fractions isolated from gliomas (*n* = 26) and non-tumor brain tissues (*n* = 6) **(Supplementary Table S1)**. Principal component analysis (PCA) revealed clear segregation of samples according to both tissue origin (tumoral or non-tumoral) and tumor IDH status along component 1, which accounted for 48% of the variance **(Fig. 1B)**. We identified 37,198 differentially methylated CpG sites (absolute Δβ > 0.2, FDR < 0.05) in myeloid cells from IDH-mutant tumors compared to those in IDH-wt gliomas **(Fig. 1C)**. Similar to the pattern observed in tumor cells enriched fractions, CD11B⁺ myeloid cells from IDH-mutant gliomas displayed a pronounced hypermethylation bias, particularly within cis-regulatory regions **(Fig. 1C)**. IDH-wt gliomas are typically enriched in MDMs, whereas the TME of IDH-mutant tumors tends to contain higher proportions of tissue-resident microglia^21–23^. However, methylation-based deconvolution^82^ of CD11B⁺ fractions revealed substantial microglial representation across tumor subtypes, despite the expected relative enrichment of MDMs in IDH-wt gliomas **(Supplementary Fig. S3)**, suggesting that the asymmetric distribution of methylome changes is unlikely to be explained solely by differences in myeloid cell composition. Consistent with this interpretation, global hypermethylation was also evident when comparing CD11B⁺ fractions from IDH-mutant gliomas to those from non-tumor brain tissues, both largely composed of microglia^21^ **(Fig. 1C)**. This later comparison revealed an even greater number of differentially methylated CpG sites and regulatory regions, with substantial overlap between the two contrasts **(Fig. 1D)**. Among the shared regions, enhancers displayed higher numbers and stronger methylation gains than promoters **(Fig. 1D)**. Enrichment analysis of genes associated with these hypermethylated regions highlighted inflammatory and immune-related pathways, including TNF-α, IFN-γ, and IL2/IL6 signaling, as well as complement and hypoxia responses **(Fig. 1E and Supplementary Table S2)**.

Transcription factors (TFs) bind to specific DNA motifs on promoters and enhancers to control cell type-specific transcriptional programs, and their affinity can be modulated by local DNA methylation^41–43^. Given that a large fraction of the hypermethylation detected in CD11B⁺ cells from IDH-mutant gliomas occurred within cis-regulatory elements across both comparisons **(Fig. 1D)**, we next analyzed the enrichment of TF-binding motifs within these regions. This analysis identified significant overrepresentation of 21 motifs in 349 hypermethylated promoters and 24 motifs in 3027 hypermethylated enhancers (p-value < 10^−12^) **(Fig. 1F)**. Among these, the *SPI1* (PU.1) motif most strongly enriched, present in 36% of enhancers and 32% of promoters. Other highly enriched motifs corresponded to a core set of TFs that coordinate environment-dependent transcriptional networks in microglia including C/EBP, AP-1, RUNX, and IRF family members, as well as NF-κB^44^. Collectively, these findings reveal extensive DNA hypermethylation within the myeloid compartment of IDH-mutant gliomas, preferentially affecting distal regulatory regions associated with immune-related circuits.

### Hypermethylation correlates with transcriptional repression of immunomodulatory programs in CD11B⁺ myeloid cells from IDH-mutant gliomas

To assess functional transcriptomic changes in the myeloid compartment of IDH-mutant tumors, we analyzed RNA-seq data from CD11B⁺ fractions isolated from gliomas (*n* = 28) and non-tumor brain tissues (*n* = 8) **(Supplementary Table S1)**. Differential expression analysis identified genes significantly upregulated or downregulated (|log₂FC| > 1, FDR < 0.05) in myeloid cells from IDH-mutant gliomas compared with either IDH-wt tumors or non-tumor brain tissues **(Supplementary Table S3)**. Functional enrichment analysis revealed that downregulated genes across both comparisons were associated with coherent inflammatory and myeloid activity programs, whereas upregulated genes showed comparatively limited functional enrichment, consistent with coordinated transcriptional repression in the IDH-mutant context **(Supplementary Table S4)**. Enriched pathways among downregulated gene sets included proinflammatory programs such as TNF-α signaling via NF-κB, INF-γ/α response, and IL6/JAK/STAT3 signaling. In addition, pathways selectively enriched in the comparison with IDH-wt gliomas were related to cell cycle regulation, proliferation, and acquisition of mesenchymal-like traits, molecular hallmarks of these tumors^24,30^ **(Supplementary Table S4)**. Together, these findings indicate that IDH-mutant tumors impose a broad repression of inflammatory and metabolic programs in the myeloid compartment.

Recent transcriptomics work showed that glioma-associated myeloid states are defined not only by lineage composition, but also by shared activity programs shaped by the tumor microenvironment^31^. To further assess functional differences across myeloid fractions, we computed single-sample gene set enrichment analysis (ssGSEA) scores for cell identity and immunomodulatory activity programs. This analysis confirmed a predominant contribution of TAM-derived expression signatures within the CD11B⁺ fractions. Microglial identity scores were comparable across tumor types, whereas macrophage and monocyte scores were higher in IDH-wt gliomas vs IDH-mutant gliomas, consistent with their greater immune infiltration **(Supplementary Fig. S4A)**. Myeloid cells from IDH-wt gliomas exhibited low activity of the microglial inflammatory program together with strong enrichment of the scavenger-associated suppressive program, which is largely absent from non-tumor brain tissue^31^. By contrast, the complement immunosuppressive program, typically excluded from hypoxic areas^31^, together with the systemic inflammatory program, were less active in myeloid cells from IDH-mutant tumors **(Supplementary Fig. S4A)**. Cell-cycle-related programs (G1-S and G2-M) displayed the highest ssGSEA scores in IDH-wt tumors, whereas the heat-shock/unfolded protein response (HS-UPR) program was significantly reduced in IDH-mutant gliomas compared to either IDH-wt or non-tumor tissue **(Supplementary Fig. S4A)**. Similarly, activation of the hypoxia program was elevated in myeloid cells from IDH-wt gliomas, whereas myeloid cells from IDH-mutant tumors exhibited trend toward repression compared with non-tumor samples. Finally, the reduced activity of shared myeloid immunomodulatory programs in IDH-mutant gliomas did not appear to be associated with tumor grade, suggesting the presence of intrinsic gene-silencing mechanisms **(Supplementary Fig. S4B)**. We next asked whether the reduced activity of these programs was accompanied by regulatory hypermethylation at their constituent genes. Consistent with this notion, a substantial fraction of representative genes within these shared myeloid activity programs were linked to hypermethylated enhancers and/or promoters in CD11B^+^ cells from IDH-mutant gliomas **(Supplementary Fig. S4C)**. This overlap was significantly greater than expected not only for the immunomodulatory activity programs but also for the hypoxia program when assessed against the full regulatory annotation background used in the methylome analysis **(Supplementary Fig. S4D)**. These genes included *CXCR4*, *CX3CR1* and *IRF8* in the microglial inflammatory program; *CD83*, *IL1B*, *CXCL8*, *NAMPT*, *OSM* and *HIF1A* in the systemic inflammatory program; and several HLA genes within the complement immunosuppressive program. Notably, additional immune mediators, including *CXCL10*, *IL18* and *CIITA*—the master transactivator of MHC class II—also exhibited regulatory hypermethylation despite not ranking among the top genes of these programs. These findings suggest that IDH-mutant-associated epigenetic priming broadly constrains shared myeloid activity programs beyond the recently defined immunomodulatory modules^31^.

To examine the extent to which DNA methylation changes may contribute to the transcriptional programs observed in CD11B⁺ cells, we intersected differentially expressed genes with genes associated with differentially methylated enhancers or promoters identified in IDH-mutant gliomas relative to IDH-wt tumors and non-tumor brain tissues. This analysis included 36 CD11B⁺ samples isolated from gliomas, 18 of which were profiled by both methylome and transcriptome analyses **(Supplementary Table S1)**. Reduced gene expression was strongly correlated with hypermethylation at cis-regulatory elements (quadrants IV), and was more prevalent at enhancers than at promoters **(Fig. 2A and Supplementary Fig. S5)**. In contrast, non-canonical associations, such as gene upregulation coupled with hypermethylation (quadrants I), were significantly underrepresented. This Integrative analysis, combining both tumoral and non-tumoral comparisons, identified 171 unique genes whose downregulated expression was linked to hypermethylated enhancers displayed by CD11B⁺ myeloid cells in IDH-mutant gliomas **(Supplementary Table S5)**.

**Figure 2.**
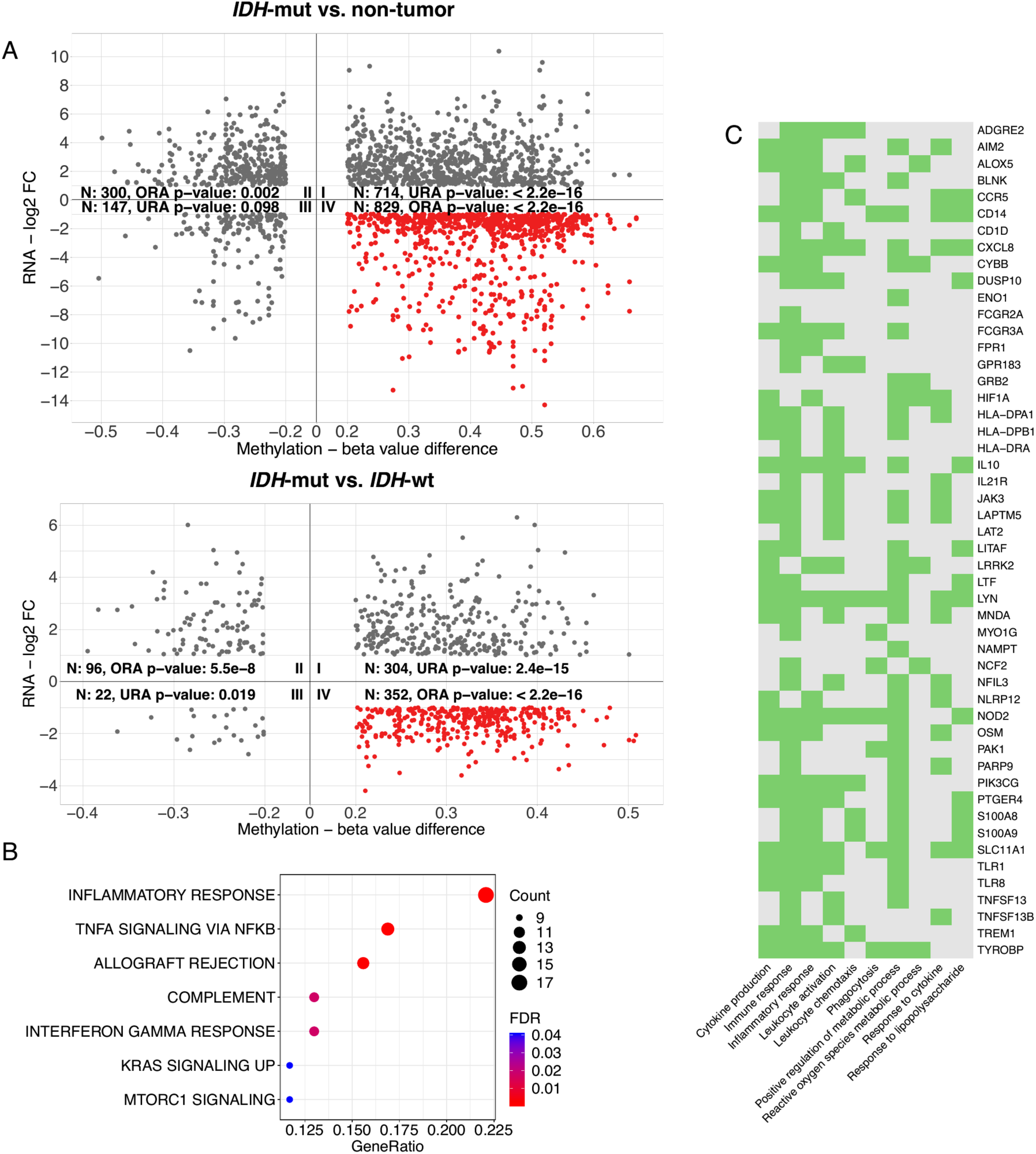
Integrated analysis of transcriptomic and methylation changes in CD11B^+^ myeloid cells from IDH-mutant gliomas. **(A)** Cartesian plots showing genes (dots) exhibiting significant changes in both expression (Y-axis) and enhancer methylation (X-axis) in CD11B^+^ fractions from IDH-mutant gliomas compared with IDH-wt tumors (upper panel) and non-tumor brain tissues (lower panel). Numbers indicate the total genes identified in each quadrant. Over-representation and under-representation of canonical methylation/expression relationships (quadrants II and IV) and non-canonical relationships (quadrants I and III) were assessed using over-representation analysis (ORA) and under-representation analysis (URA), respectively. Quadrant IV (highlighted in red) contains genes exhibiting enhancer hypermethylation together with transcriptional downregulation. **(B)** Bubble plot showing Hallmark gene sets enriched within the integrative gene set (*n* = 171) generated by combining genes from quadrant IV across both comparison settings. Dot size and color code indicate the number of genes contributing to each enriched category and the corresponding FDR value, respectively (FDR < 0.05). **(C)** Plot showing a subset of genes from the integrative gene set enriched in Gene Ontology biological processes related to immune response and metabolism. Green boxes indicate the presence of enhancer hypermethylation associated with each gene.

The integrated gene list included regulators of TNF-α signaling via NF-κB, complement activation, IFN-γ response, and mTORC1 signaling pathways **(Fig. 2B)**. Enriched biological processes included cellular response to stimulus, cytokine production, leukocyte activation, and positive regulation of metabolism **(Fig. 2C)**. Affected genes in these categories comprised proinflammatory cytokines (*CXCL8, OSM*), TNF ligand superfamily members (*TNFSF13, TNFS13B*), the anti-inflammatory cytokine *IL10*, chemokine receptors (*CCR5, FPR1, TLR1/8, IL21R*), and *HLA* genes, together with regulators of glycolysis/hypoxia (*HIF1A, CD44*). Additional components included mediators of immune activation (*TYROBP)*, NAD^+^ biosynthesis (*NAMPT*), and oxidative metabolism (*CYBB*) **(Fig. 2C)**. While *CXCL10, IL18* and *CIITA*, were selectively repressed in comparison with IDH-wt tumors, *IL1B* and *IL4R* were specifically downregulated relative to non-tumor brain tissues, indicating both shared and context-dependent repression patterns.

To validate these findings at the protein level, we performed targeted proteomic profiling of inflammation- and immune-related proteins in an independent series of CD11B⁺ fractions isolated from gliomas (*n* = 13) and from non-tumor brain tissues (*n* = 3) **(Supplementary Table S1)**. Across the full panel, myeloid cells from IDH-mutant tumors exhibited a pronounced global decrease in protein abundance relative to non-tumor samples **(Supplementary Fig. S6A and Supplementary Table S6)**. In comparison with IDH-wt tumors, protein changes were directionally consistent but of smaller magnitude **(Supplementary Fig. S6B and Supplementary Table S6)**. Protein-level differences between the two comparisons were moderately but significantly correlated (Spearman ρ and Pearson r ≈ 0.5; both p < 10⁻⁷⁸), indicating a coherent IDH mutation-associated proteomic program in myeloid cells. Of the 1,034 proteins profiled, 101 corresponded to genes identified in the integrative methylation-transcriptome analysis from either comparison **(Fig. 2A, quadrant IV)**. Notably, this protein subset exhibited a significantly stronger negative shift in ΔNPX (change in normalized protein expression) than the full panel, supporting convergence between enhancer hypermethylation, transcriptional repression, and reduced protein abundance **(Supplementary Fig. S6A and S6B)**. A core group of 17 proteins derived from genes shared across both integrative comparisons showed consistent downregulation in both contrasts **(Supplementary Fig. S6C and Supplementary Table S6)**. These included inflammatory mediators such as CXCL8, OSM, TNFSF13, TLR1, and HLA-A. In contrast, HIF1A and CXCL10 repression was restricted to the transcriptomic level, whereas the suppressive (”M2-like”) factors ARG1 and TGFBR1 were decreased at both mRNA and protein levels **(Supplementary Fig. S6C)** despite lacking enhancer hypermethylation, suggesting additional regulatory mechanisms. Together, these findings indicate that enhancer hypermethylation is associated with suppression of inflammatory and immune signaling, suggesting a globally hyporesponsive state in myeloid cells from IDH-mutant gliomas.

### Enhancer hypermethylation in CD11B^+^ cells is linked to reduced activity of core microglial transcription factors

To assess how enhancer hypermethylation relates to transcriptional regulation at the loci highlighted in quadrants I and IV **(Fig. 2A)**, we inferred transcription factor activity from RNA-seq data using high-confidence TF-gene interactions (*n* = 1,209 human TFs)^45,46^. Regulon analysis identified 45 TFs with differential activity (FDR < 0.05; Wilcoxon’s test) according to tumor IDH status, with most showing reduced inferred activity in IDH-mutant tumors **(Fig. 3A)**. Consistent with the motifs enriched in hypermethylated enhancers **(Fig. 1F)**, we found that lineage-determining and stimulus-responsive microglial TFs were among the most affected. These included *SPI1* (PU.1), the AP-1 complex (*ATF2, FOS, JUN*), and *CEBPB*, all previously linked to DNA methylation-dependent regulation^47–49^. Additional TFs with reduced inferred activity, such as *NFATC2,* subunit *RELA/p65* (NF-κB), and STAT3 are known to occupy microglial super-enhancers^44^. In contrast, only a small subset of TFs appeared more active in myeloid cells from IDH-mutant gliomas, including *TP63, TP73, SNAI1, CTNNB1, EPAS1/HIF2A*, and *KLF4*, for which methylated CpGs may instead facilitate binding^50^. TF-target gene regulatory network analysis further indicated that genes controlled by core microglial TFs involved in inflammation and antigen presentation were downregulated, consistent with disruption of cooperative TF activity **(Fig. 3B)**. Collectively, these results identify enhancers as major targets of tumor-associated microenvironmental perturbation and support a link between DNA hypermethylation, reduced TF activity, and the blunted activation state of myeloid cells in IDH-mutant gliomas.

**Figure 3.**
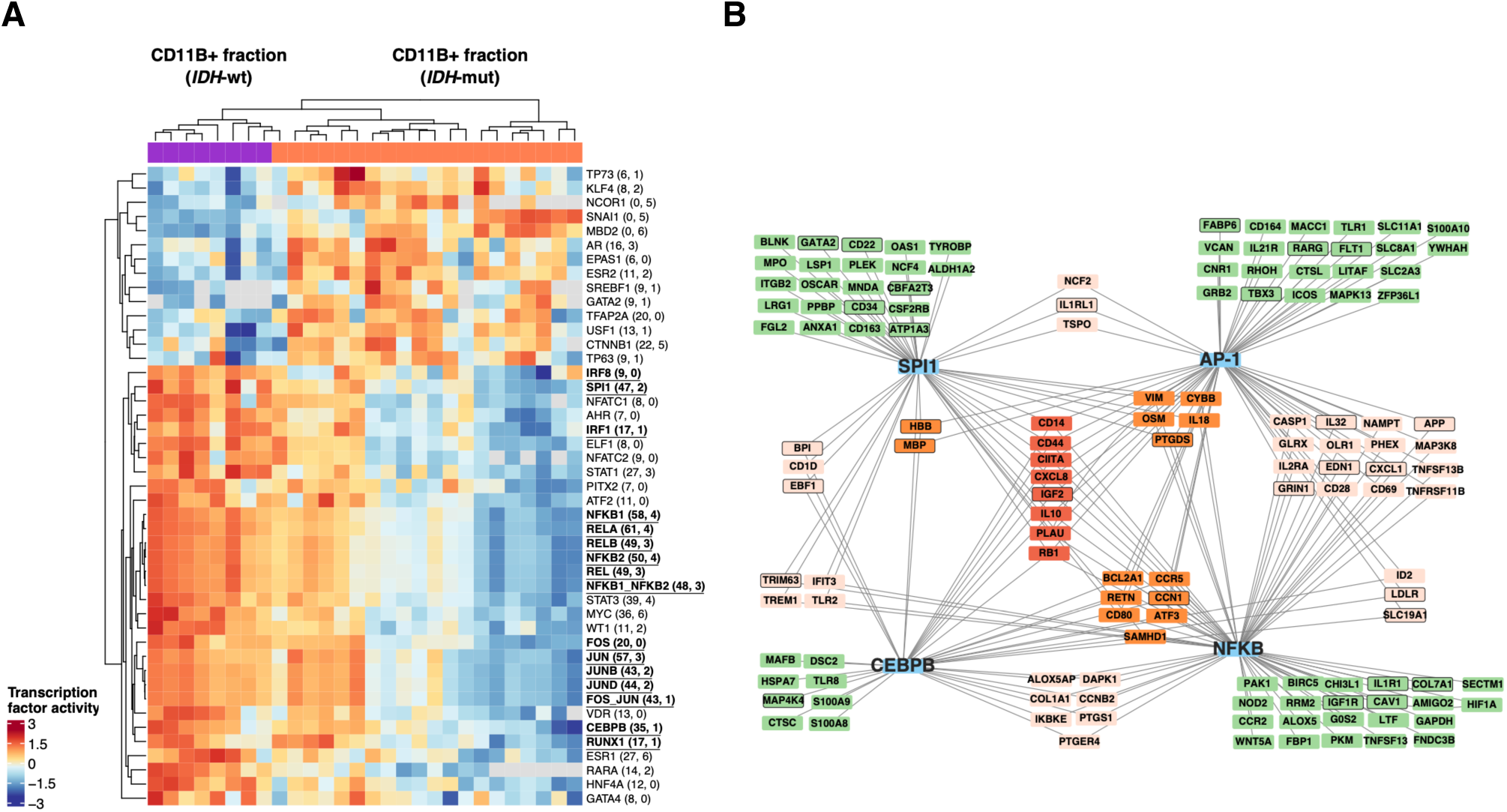
Regulon analysis reveals reduced activity of microglial TFs associated with enhancer hypermethylation in CD11B^+^ cells from IDH-mutant gliomas. **(A)** Heatmap showing TFs exhibiting differential activity in CD11B^+^ fractions according to the IDH status of the corresponding tumors. TF target genes were retrieved from the CollecTRI database and intersected with genes displaying differential expression together with enhancer hypermethylation in myeloid cells from IDH-mutant gliomas (quadrants I and IV, Fig. 2A). Numbers in brackets indicate the number of positively and negatively regulated targets associated with each TF. Core microglial TFs identified in the previous motif enrichment analysis are shown in bold and underlined. Unsupervised clustering was performed using TF activity z-scores, where red and blue indicate high and low TF activity, respectively, and gray indicates non-significant activity. **(B)** Gene regulatory network of four representative TFs (blue nodes) exhibiting reduced activity in CD11B^+^ cells from IDH-mutant relative to IDH-wt gliomas. TF target genes are colored according to the number of TFs regulating them (red, four TFs; orange, three TFs; pink, two TFs; green, one TF). Targets outlined in black correspond to genes from quadrant IV (hypermethylated enhancers associated with downregulated expression), whereas non-outlined targets correspond to genes from quadrant I (hypermethylated enhancers associated with upregulated expression).

### D-2HG reduces TET activity and impairs DNA methylation turnover at selected enhancers in primary human microglia

We hypothesized that D-2HG, the oncometabolite produced by IDH-mutant glioma cells and released into the TME, alters the DNA methylation landscape of myeloid cells. Ten-eleven translocation (TET) enzymes are α-KG-dependent dioxygenases that catalyze the oxidation of 5-methylcytosine (5mC) to 5-hydroxymethylcytosine (5hmC)^51^, the first step of active DNA demethylation **(Fig. 4A)**. Inhibition of this process by D-2HG reduces 5hmC levels and leads to progressive accumulation of 5mC over successive cell divisions, ultimately establishing the G-CIMP-high phenotype characteristic of IDH-mutant gliomas^52,53^. To determine whether a similar mechanism operates within the myeloid compartment, we quantified global 5mC and 5hmC levels, and applied epigenetic clock estimates to independent CD11B⁺ fractions isolated from IDH-mutant gliomas (*n* = 4) and non-tumor brain tissues (*n* = 5) **(Supplementary Table S1)**. As expected, CD11B⁻ tumor fractions from IDH-mutant gliomas displayed markedly elevated 5mC/5hmC ratios compared with non-tumor samples **(Fig. 4B)**. Importantly, a significant increase in this ratio was also observed in the corresponding CD11B⁺ myeloid fractions and was accompanied by increased epigenetic mitotic age estimates relative to non-tumor brain tissues **(Fig. 4C and 4D)**. Given that both sample groups are largely enriched in microglia, these findings are unlikely to be explained solely by differences in myeloid composition and instead support chronic epigenetic remodeling within the resident microglial population. Together, these findings are consistent with impaired TET-mediated demethylation associated with D-2HG exposure, contributing to the global DNA hypermethylation observed in myeloid cells from IDH-mutant gliomas.

**Figure 4.**
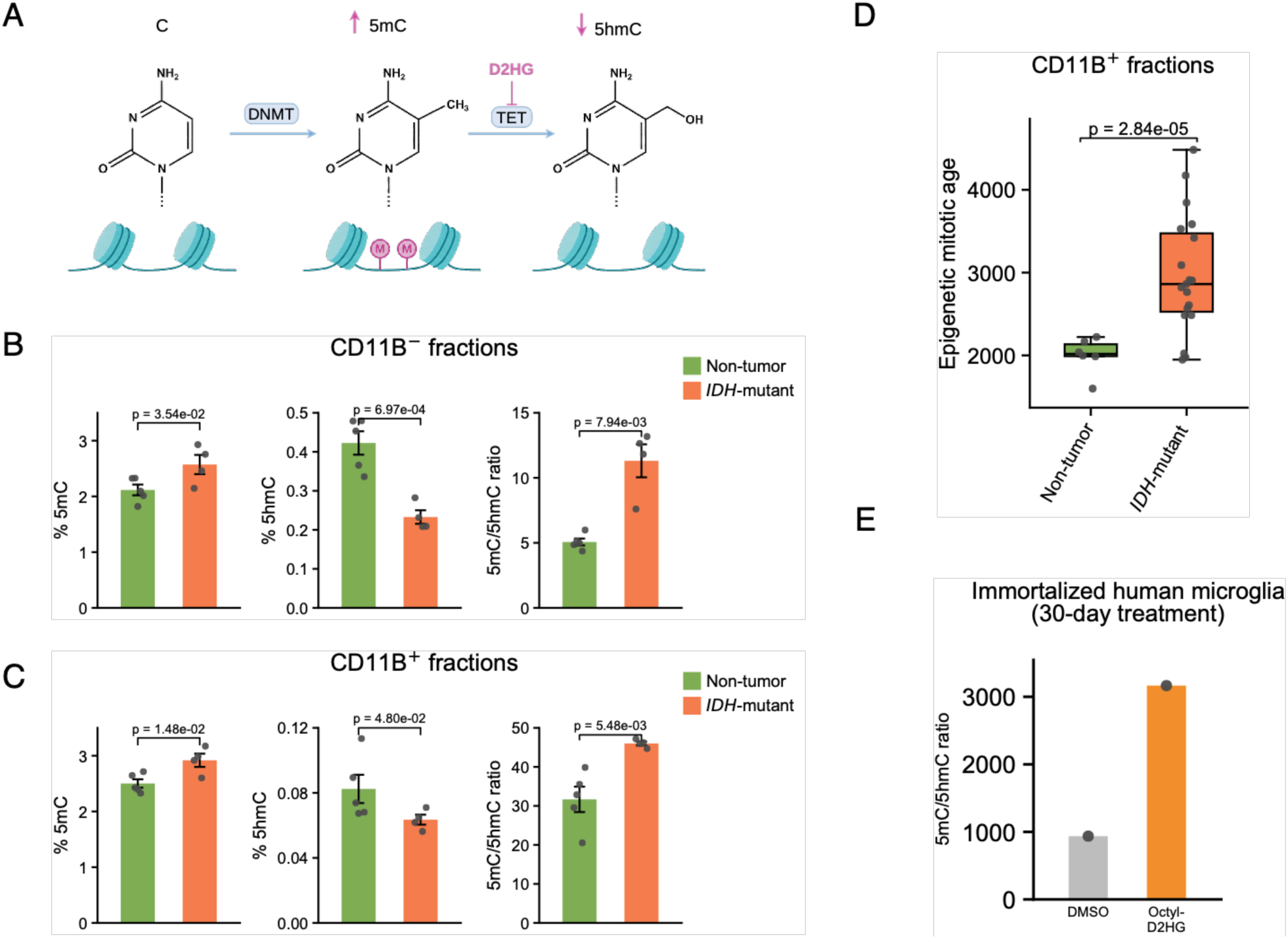
Elevated 5mC/5hmC ratio and epigenetic mitotic age in CD11B^+^ myeloid cells from IDH-mutant gliomas. **(A)** Schematic representation of α-KG-dependent oxidation of 5mC by TET enzymes and competitive inhibition of this process by D-2HG in IDH-mutant gliomas. **(B-C)** Global levels of 5mC and 5hmC, and corresponding 5mC/5hmC ratios, measured by LC-MS/MS in CD11B⁻ (upper panels) and CD11B⁺ (lower panels) fractions from IDH-mutant gliomas (*n* = 4) and non-tumor brain tissues (*n* = 5), which served as controls. Data are presented as mean ± SEM. P-values were calculated using one-sided Welch’s tests, except for the 5mC/5hmC ratio in the CD11B⁻ fraction (one-sided Wilcoxon’s test). **(D)** Epigenetic mitotic age analysis performed using the epiTOC2 model to estimate age-adjusted replication-associated methylation accumulation within the CD11B^+^ myeloid compartment of IDH-mutant tumors and non-tumor brain tissues. P-values were calculated using Welch’s test. **(E)** Global 5mC/5hmC ratios measured by LC-MS/MS in immortalized human microglia following prolonged exposure to octyl-D-2HG or DMSO. Data represent a single long-term treatment experiment.

Because tissue-resident microglia are chronically exposed to high concentrations of D-2HG from tumor initiation onward, we asked whether this prolonged exposure preferentially alters their DNA methylome, in contrast to infiltrating immune cells, which encounter only transient exposure and in which prior studies have suggested predominantly non-epigenetic effects^54^. To test this hypothesis directly, we first examined whether prolonged D-2HG exposure alters global DNA methylation in immortalized human microglia. Consistent with reduced TET activity, prolonged treatment with cell-permeable octyl-D-2HG increased the 5mC/5hmC ratio relative to DMSO-treated controls after 30 days of exposure **(Fig. 4E)**. However, as commonly observed in immortalized systems, baseline 5hmC levels were very low, limiting the extent of detectable demethylation changes. We therefore established primary cultures of human microglia from tissue aspirates obtained during surgery for glioma or drug-resistant epilepsy to assess D-2HG effects in a more physiologically relevant setting **(Supplementary Fig. S7A)**. Cells maintained in serum-free, defined medium^55^ remained viable for up to 15 days, with >95% expressing the microglial markers IBA1 and TMEM119 **(Supplementary Fig. S7B)** and <5% showing proliferation (KI67^+^; data not shown). These cultures preserved a stable microglial transcriptional signature by RNA-seq and remained functionally responsive, exhibiting a robust proinflammatory response to lipopolysaccharide (LPS) stimulation at 48 h or 14 days **(Supplementary Fig. S7C and S7D)**.

To determine whether human microglia can internalize D-2HG from their environment, we measured intracellular levels after 48 hours of exposure to a concentration comparable to that estimated in the TME of IDH-mutant gliomas^5,6^, which did not affect cell viability. Intracellular D-2HG levels measured by fluorometric assay in treated microglia reached values similar to those observed in *bona fide* IDH-mutant cells, and LC-MS/MS confirmed intracellular accumulation of 2HG in independently treated primary microglial cultures **(Fig. 5A)**. This uptake was sufficient to significantly inhibit TET enzymatic activity **(Fig. 5B)**. We then exposed primary microglial cultures to D-2HG for a prolonged period and observed a progressive increase in global 5hmC levels in controls, whereas 5hmC remained stable in D-2HG-treated cells, resulting in a significant reduction at 14 days **(Fig. 5C)**. However, in contrast to immortalized microglia, no concomitant increase in 5mC levels was detected (data not shown), consistent with the limited proliferative capacity of the primary culture system. We next asked whether D-2HG might nevertheless affect TET-mediated demethylation dynamics at specific genomic elements, particularly enhancers, which are characterized by high methylation turnover^56,57^. To address this, we used a reduced representation methylation sequencing approach^58,59^ that enables quantitative base-resolution mapping of 5mC and 5hmC across entire enhancer regions in primary human microglia derived from three non-tumor brain tissues following 14 days of D-2HG exposure. We specifically examined DNA methylation dynamics at TF-binding motifs embedeed within enhancers identified as hypermethylated in CD11B⁺ cells from IDH-mutant gliomas, as methylation at these sites may influence TF occupancy **(Fig. 5D)**.

**Figure 5.**
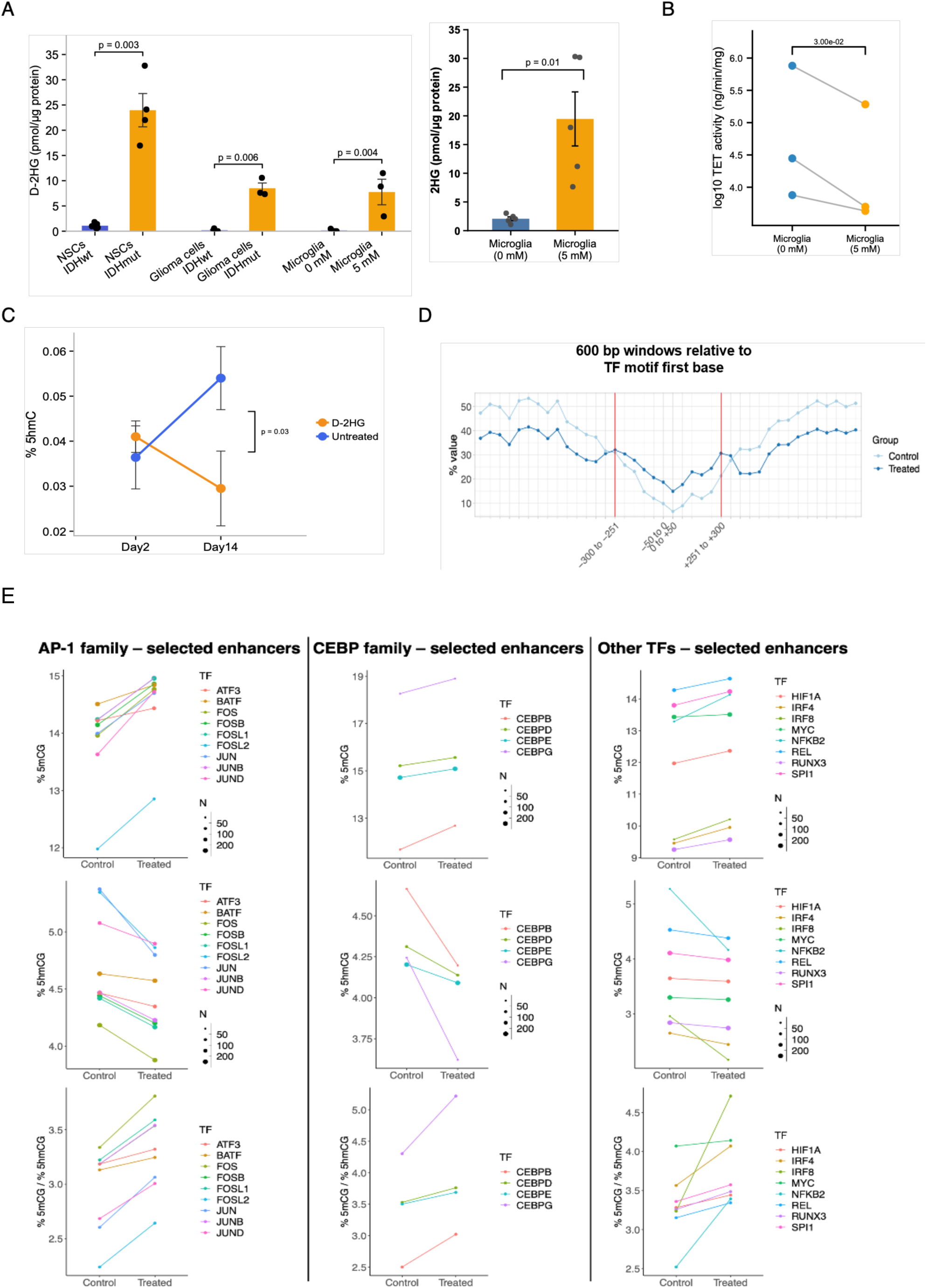
D-2HG impairs TET-mediated demethylation turnover at specific enhancers in primary human microglia. **(A)** D-2HG levels were determined using a fluorometric assay in cell pellets from cKI neural stem cells (NSCs; IDH-wt, *n* = 5; IDH-mutant, *n* = 4), patient-derived glioma cells (IDH-wt, *n* = 4; IDH-mutant, *n* = 3), and primary human microglia (*n* = 3) treated with D-2HG (5 mM) for 48 h. Intracellular 2HG levels were further quantified by LC-MS in independently treated primary human microglial cultures (*n* = 5). Metabolite levels were normalized to total protein content. Data are presented as mean ± SEM. P-values were calculated using a one-sided paired Wilcoxon signed-rank test. **(B)** TET enzymatic activity, measured as hydroxymethylated product formation (ng), after 1 hour incubation of substrate with protein extracts from untreated or D-2HG-treated (5 mM, 48 h) primary microglial cultures (*n* = 3). P-value was calculated using a one-sided paired t-test. **(C)** Global 5hmC levels in primary microglial cultures (*n* = 5), determined by LC-MS/MS following 48 h or 14 days of treatment with D-2HG (5 mM). Data are presented as mean ± SEM. P-value was calculated using a paired Wilcoxon signed-rank test. **(D)** Schematic representation of the sequencing-based analysis used to quantify 5mC and 5hmC levels within defined windows surrounding TF-binding motifs located in enhancers from primary microglia treated with D-2HG for 14 days and paired untreated controls. **(E)** Plots showing separate analyses of 5mC and 5hmC percentages (upper panels) surrounding binding motifs for the indicated TFs in D-2HG-treated microglia and paired untreated controls (*n* = 3). Only enhancers shared across samples were included in the analysis. Dot size indicates the number of enhancers analyzed in a paired manner between conditions. Corresponding 5mC/5hmC ratios (lower panels), are also shown. TF-binding motifs included in this analysis were located within enhancers exhibiting hypermethylation in CD11B^+^ fractions from IDH-mutant gliomas relative to non-tumor brain tissues in the *ex vivo* cohort (Fig. 1C).

We detected a modest but reproducible increase in 5mC/5hmC ratios around TF-binding motifs belonging to the AP-1, C/EBP, and NF-κB families in D-2HG-treated primary microglia compared with paired untreated controls **(Fig. 5E)**. Similar patterns were obtained for motifs bound by *SPI1* (PU.1), *IRF4/8*, *RUNX3*, *HIF1A*, and *MYC*, which were likewise implicated in the *ex vivo* analyses. In sharp contrast, 5mC and 5hmC levels exhibited opposite trends when the analysis was performed across the full set of covered enhancers, used as background reference **(Supplementary Fig. S8A)**. Importantly, genes associated with enhancers displaying both hypermethylation and hypohydroxymethylation **(Supplementary Table S7)** were significantly enriched among downregulated transcripts, consistent with D-2HG-induced enhancer hypermethylation contributing to repression of activation-related genes **(Supplementary Fig. S8B and S8C)**.

To further assess whether prolonged D-2HG exposure induces similar enhancer remodeling *in vivo*, we established a xenograft model in which D-2HG was continuously delivered to the tumor microenvironment using osmotic pumps. CD11B⁺ myeloid fractions isolated from treated mice exhibited reduced 5hmC levels at orthologous enhancers enriched for TF-binding motifs previously identified in the *ex vivo* analyses, whereas opposite trends were observed across the full set of covered enhancers used as background reference broadly recapitulating the alterations observed in primary human microglia exposed to D-2HG *in vitro* **(Supplementary Fig. S9)**. Together, these findings indicate that D-2HG preferentially alters DNA methylation turnover at a subset of microglial enhancers, thereby reshaping an epigenetic landscape associated with activation responses.

### D-2HG induces a hyporesponsive transcriptional state in LPS-stimulated microglia that is partially reversed following IDH inhibition in patients

The enhancers affected by D-2HG treatment in our *in vitro* experiments are likely to be transcriptionally engaged upon macrophage stimulation^60^. We therefore hypothesized that D-2HG-induced methylome priming could impair the activation of enhancer elements that regulate the proinflammatory capacity of human microglia. To address this, we evaluated transcriptional changes in microglial cultures pretreated with D-2HG for 14 days and subsequently challenged with LPS, a TLR4 agonist for 6 hours **(Fig. 6A)**. Gene set enrichment analysis (GSEA) of paired samples revealed that D-2HG treatment consistently attenuated LPS-induced microglial activation **(Fig. 6B)**.

**Figure 6.**
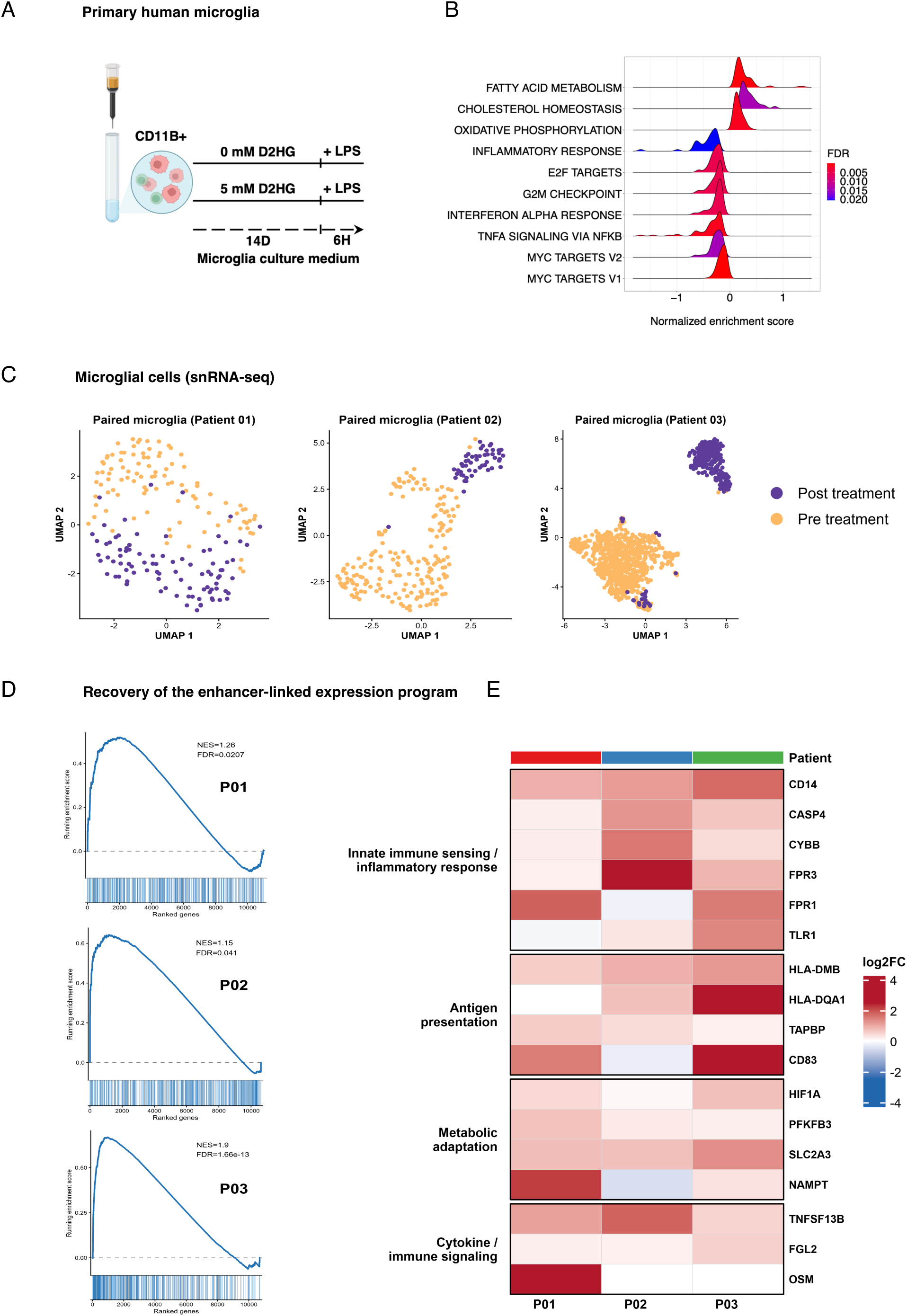
D-2HG represses transcriptional expression of microglial activation-related genes. **(A)** Experimental workflow for the LPS challenge assay in primary human microglial cultures (*n* = 4) pre-treated with D-2HG for 14 days. Transcriptomic analyses (RNA-seq) were performed using a patient-paired design. **(B)** GSEA ridge plot showing Hallmark gene sets enriched in D-2HG-treated microglia 6 h after LPS stimulation. Positive and negative normalized enrichment scores (NES) correspond to gene sets enriched among upregulated and downregulated genes, respectively (FDR < 0.05). **(C)** Uniform Manifold Approximation and Projection (UMAP) plots of microglial cells from three patients analyzed by snRNA-seq before and after treatment with mutant IDH inhibitors. **(D)** Preranked gene-set enrichment analysis performed on the post- versus pre-treatment differential expression signature of microglia from each patient. Genes were ranked using a signed differential expression metric, with genes induced after treatment placed at the top of the ranked list and downregulated genes at the bottom. The queried gene set corresponded to the core enhancer-linked repression program, defined as genes associated with enhancer hypermethylation together with transcriptional downregulation in CD11B⁺ fractions of IDH-mutant gliomas in the *ex vivo* analyses. Running enrichment curves show the distribution of this gene set across each ranked signature. NES and FDR are indicated. **(E)** Heatmap showing avg_log2FC values for representative genes induced after treatment with mutant IDH inhibitors in microglia. Genes were selected as biologically informative components of the enhancer-linked repression program based on convergent evidence from independent epigenetic, transcriptomic, and functional analyses.

Specifically, D-2HG-treated cells exhibited a hyporesponsive state characterized by enhanced OXPHOS and fatty acid metabolism together with suppression of inflammatory response pathways, including TNF-α signaling via NF-κB **(Fig. 6B)**. To validate these findings, we first assessed the expression of representative LPS-responsive cytokines in independent primary microglial cultures. RT-qPCR analysis confirmed reduced induction of inflammatory mediators *TNF* and *IL6* following prolonged D-2HG exposure **(Supplementary Fig. S10A)**. We next performed targeted proteomic profiling, as in the *ex vivo* setting, analyzing cell lysates from human immortalized microglia and supernatants from primary microglial cultures **(Supplementary Fig. S10B)**. Although the effect size was modest, D-2HG also attenuated the expression of LPS-induced proteins in both systems. Notably several of these proteins were also repressed in CD11B^+^ fractions from IDH-mutant tumors compared with non-tumor brain tissues **(Supplementary Table S6)**. Assessment of mitochondrial respiration further demonstrated increased basal and maximal respiration in D-2HG-treated microglia, even after LPS stimulation **(Supplementary Fig. S10C)**, consistent with altered metabolic adaptation during microglial activation.

Together, these *in vitro* epigenomic, transcriptomic and functional data recapitulate the *ex vivo* hyporesponsive phenotype, supporting a role for D-2HG in constraining the deployment of microglial immune programs through altered methylation turnover at enhancer elements.

Finally, to assess whether the microglial hyporesponsive state associated with IDH-mutant gliomas could be modulated by IDH-targeted therapy, we analyzed snRNA-seq data from three paired tumor samples collected before and after mutant IDH inhibitor exposure in patients who did not corticosteroids **(Supplementary Table S1)**^61^. Myeloid cells in these samples corresponded predominantly to microglia rather than MDMs, based on canonical marker expression **(Supplementary Fig. S11)**. Across patients, on-treatment microglia exhibited transcriptional changes compared with pre-treatment microglia **(Fig. 6C and Supplementary Table S8)**.

To determine whether these transcriptional changes were related to the enhancer-linked regulatory program identified in CD11B⁺ cells from IDH-mutant gliomas, we performed preranked GSEA using the full post- versus pre-treatment microglial differential-expression signature from each patient. Genes linked to the recurrent set of hypermethylated enhancers associated with reduced target-gene expression **(Fig. 2A, quadrants IV)** were significantly enriched toward the post-treatment upregulated end of the ranked signatures (FDR < 0.05) **(Fig. 6D)**. Analysis of these transcriptional changes revealed consistent re-expression of a subset of genes whose downregulation corresponded to hypermethylated enhancers **(Fig. 2A, quadrants IV)** across all patients treated with mutant IDH inhibitors, alongside variability in the magnitude and composition of expression changes between tumors. Re-expressed genes span innate immune sensing, antigen presentation, metabolic adaptation, and immune signaling **(Fig. 6E)**. Together, these results indicate that IDH inhibition restores expression of microglial immune-related programs linked to an enhancer-associated regulatory network epigenetically constrained in IDH-mutant gliomas.

## Discussion

Understanding how the glioma microenvironment shapes the transcriptional programs and functional states of TAMs is central to developing new therapeutic strategies. Prior studies have shown that tumor IDH status defines the transcriptomic profile of these cells^21–24,30^, reflecting distinct metabolic and signaling contexts. More recently, enhancer dynamics and the coordinated activity of lineage-determining and stimulus-responsive TFs^62,63^ have emerged as important determinants of glioma-associated myeloid states^31^. However, the contribution of DNA methylation to these regulatory circuits, and its modulation by tumor-derived metabolites, has remained largely unexplored. Here, we provide evidence that chronic exposure of tissue-resident microglia to the oncometabolite D-2HG impairs TET-dependent demethylation turnover, remodels the enhancer landscape, and promotes a hyporesponsive immune state. This form of epigenetic priming represents an early mechanism of immune evasion, likely contributing to the immunologically cold microenvironment characteristic of IDH-mutant gliomas.

Our *ex vivo* profiling of CD11B⁺ myeloid fractions from gliomas revealed a marked shift toward DNA hypermethylation in IDH-mutant tumors. Importantly, a significant subset of hypermethylated cis-regulatory regions was shared when CD11B⁺ cells from IDH-mutant gliomas were compared with both IDH-wt tumors and non-tumor brain tissues. Because these reference groups differ in their relative contributions of MDMs and resident microglia^21–23^, the convergence of these regions argues against a purely compositional explanation and supports the presence of genuine epigenetic changes within the myeloid compartment. Furthermore, microglial identity scores were consistently high across the different groups, pointing to functional differences within this resident myeloid population. Together, these findings indicate that the widespread hypermethylation detected in the myeloid compartment of IDH-mutant gliomas primarily reflects epigenetic reprogramming of tissue-resident microglia rather than differences in MDM infiltration alone, and reinforce the concept that the TME actively educates tissue-resident myeloid cells^64,65^.

Enhancer elements, which encode microglial identity and mediate responses to environmental cues, were the principal targets of DNA hypermethylation, with a substantial subset linked to reduced expression of their target genes. The preferential impact on enhancers rather than proximal promoters is consistent with their higher methylation turnover and sensitivity to TET2 inhibition^55,66^. CpG methylation within or flanking TF motifs can influence DNA binding by altering local DNA conformation and nucleosome positioning—sometimes enhancing, but more often attenuating TF occupancy^67–69^. Accordingly, our regulon analysis revealed reduced activity of core microglial TFs, including the pioneer factor PU.1 and the stimulus-responsive NF-κB and AP-1 families, reflected in the coordinated repression of inflammatory, antigen-presenting, and glycolytic genes. Recent work has highlighted dynamic enhancer activity as a critical determinant of microglial plasticity^62,63^. Based on complementary analyses of human microglia purified from gliomas or cultured in the presence of D-2HG, our findings provide evidence that distal enhancers can become epigenetically altered in a tumoral context, potentially through oncometabolite signaling in the IDH-mutant TME.

Placing these observations in the broader framework of glioma-associated myeloid biology, our work provides a mechanistic basis for the contrasting immunomodulatory programs of IDH-mutant and IDH-wt gliomas, each shaped by distinct microenvironmental pressures. In the IDH-wt context, hypoxia and scavenger programs are highly active, likely driven by HIF1α stabilization^70^ together with strong AP-1 activity, generating metabolically active TAM states associated with suppressive and tissue-remodeling features^30–32^. Whether NF-κB/TET2-mediated demethylation described in inflammatory MDMs cultured under hypoxic conditions^71^ also contributes to these programs in IDH-wt gliomas remains unknown, particularly since DNA methylation dynamics were not assessed in prior transcriptome-based analyses^30,31^.

By contrast, the microenvironment of IDH-mutant gliomas appears to impose an epigenetic form of control whereby enhancer hypermethylation limits the NF-κB-driven systemic inflammatory and complement-associated myeloid activity programs, the latter typically excluded from hypoxic niches. Moreover, we observed hypermethylation linked to downregulation of the glycolytic master regulator *HIF1A* and other hypoxia-responsive genes previously reported to be underexpressed in bulk IDH-mutant tumors and cells under hypoxic culture conditions^72^. Thus, enhancer hypermethylation seems to raise the threshold for microglial activation, limiting their ability to engage in inflammatory signaling and metabolic adaptation even when appropriate stimuli are present. This persistent epigenetic restriction, shaped in part by the oncometabolite D-2HG, provides a potential mechanistic explanation for the reduced expression of inflammatory cytokines and chemokines previously reported in IDH-mutant gliomas, helping reconcile why these tumors harbor abundant microglia yet remain immunologically quiet^73,74^.

Microglia are long-lived CNS-resident cells and may undergo proliferation in response to stimuli. In IDH-mutant gliomas, these cells are chronically exposed to D-2HG, making them particularly susceptible to progressive enhancer hypermethylation and epigenetic reprogramming. By combining short- and long-term D-2HG exposure paradigms in primary and immortalized human microglia, respectively, we demonstrated that D-2HG inhibits 5mC oxidation in non-proliferating cells whereas it drives global DNA hypermethylation ove time only in immortalized cells. However, single-base– resolution profiling in primary human microglia as well as in CD11B^+^ cells from the *in vivo* D-2HG delivery model showed that this oncometabolite readily shifts the 5mC/5hmC balance at enhancers found to be hypermethylated *ex vivo,* thus supporting impairment of TET-mediated demethylation at these regulatory regions^56,57^. Functionally, D-2HG–treated primary microglia showed blunted induction of cytokine and NF-κB–driven transcriptional programs after LPS stimulation and maintained an OXPHOS-biased metabolic state, further suggesting impaired metabolic adaptation during activation^75,76^.

Considering the time-dependent effects of D-2HG on the microglial methylome, our findings support a model in which DNA hypermethylation at enhancer elements is associated with reduced activity of core microglial TFs. This process may impair the ability of microglia to mount proinflammatory responses and constrain activation-associated metabolic adaptation. However, because D-2HG likely affects multiple α-KG-dependent dioxygenases, including histone demethylases, altered chromatin remodeling may also contribute to the observed phenotype. Our model also does not exclude contributions from non-epigenetic mechanisms, including signaling pathways previously shown to be targeted by D-2HG in other cell types^54^. In particular, acute octyl-D-2HG treatment has been reported to dampen LPS-induced cytokine expression in immortalized mouse microglial cells through an AMP-activated protein kinase (AMPK)-mTOR-dependent reduction of NF-κB activity^77^. Nonetheless, the overall concordance between our *ex vivo*, *in vitro*, and *in vivo* findings provides strong evidence that chronic D-2HG exposure remodels microglial enhancer landscapes, thereby contributing to their altered functional state in IDH-mutant gliomas.

In line with this model, analysis of microglia from IDH-mutant patients treated with the mutant IDH inhibitors—known to deplete D-2HG levels—revealed re-expression of immune-related genes associated with enhancers hypermethylated in myeloid cells from IDH-mutant gliomas. This in-human observation supports the notion that D-2HG acts as a suppressive metabolite contributing to the epigenetic silencing of microglial immune responses, and that relieving this D-2HG-dependent brake can re-engage these circuits. Such reactivation may help shift the immunologically cold microenvironment toward a state more permissive to antitumor immunity and sensitizing it to subsequent immune interventions^78,79^. Whether this reprogramming is complete, partial, or dependent on exposure duration remains to be determined, particularly given that enhancer hypermethylation can be relatively stable once established.

### Conclusions and limitations

In sum, our findings demonstrate that the oncometabolite D-2HG drives enhancer-level epigenetic silencing of microglial activation, defining a reversible component of tumor-associated immune evasion. By extending the impact of IDH inhibition beyond tumor-intrinsic effects to encompass reprogramming of the myeloid compartment, this work provides a mechanistic framework for interpreting both biological effects and clinical responses to IDH-targeted therapy and for informing rational combinations with immune-modulating treatments for patients with IDH-mutant gliomas^80^. Nevertheless, certain limitations should guide interpretation. Our *ex vivo* analyses rely on bulk profiling of CD11B⁺ cells, although complementary deconvolution analyses, primary human microglia, and *in vivo* validation support a predominant contribution of tissue-resident microglia.

Future single-cell epigenomic approaches integrating enhancer methylation, chromatin accessibility, and transcriptional state will help refine cell-type attribution and regulatory circuitry. We also did not directly assess local chromatin remodeling at targeted enhancers, which may influence accessibility alongside DNA methylation.

## Author’s Contributions

**A.L.:** Conceptualization, methodology, investigation, formal analysis, data curation, writing–review and editing. **P.P.:** Conceptualization, formal analysis, data curation, visualization, writing–review and editing. **S.S.:** Data curation, formal analysis, and investigation. **B.M.:** Resources and methodology. **L.D.:** Data curation, formal analysis, visualization, writing–review and editing. **Q.R. S.J.:** Data curation and investigation. **C.B.:** Formal analysis, investigation, and visualization. **K.S.:** Methodology, writing–review and editing. **L.S., Y.H., N.P.:** Data curation, formal analysis, and investigation. **P.M.:** Resources. **K.L., A.A.:** Data curation and bioinformatics support. **M.V., A.I., C.B.:** Resources. **A.D., I.F.:** Investigation. **P.J.K., L.J.M.:** Investigation and formal analysis. **K.M., S.T.,** Resources and data curation. **E.H.:** Resources. **E.D. and E.E.-H.:** Investigation, formal analysis, visualization, writing–review and editing. **J.B., L.A.S.:** Data curation, formal analysis, writing–review and editing. **M.L.S., M.C., A.I., G.H.:** Data curation, resources, writing–review and editing. **G.A.K.:** Investigation, formal analysis, data interpretation writing–review and editing. **F.B., M.T.:** Resources, funding acquisition, writing-review and editing. **M.M.:** Conceptualization, methodology, formal analysis, data interpretation, writing–review and editing. **M.S.:** Conceptualization, formal analysis, data interpretation, supervision, funding acquisition, writing– review and editing. **L.J.C.-V.:** Conceptualization, methodology, formal analysis, data interpretation, visualization, project administration, supervision, funding acquisition, and writing–original draft.

## Acknowledgements

The authors express their deepest gratitude to Prof. Dr. Laurent Capelle for his insightful feedback and invaluable contributions to the collection of neurosurgical specimens for research purposes, including the present study. The authors also thank the personnel of the iGenSeq (sequencing), Celis (cell culture), and Histomics (histology) platforms at the ICM for their technical support, as well as Stéphanie Jouannet, Julie Jardon, Coralie Gimonnet, and Giovanni Scala for assistance with experiments and bioinformatics analyses. This work was supported by the Ligue Nationale contre le Cancer and by the following grants: INCa-DGOS-Inserm 12560; Site de Recherche Intégré sur le Cancer (SiRIC CURAMUS); French National Cancer Institute (INCa-PLBIO22-243); Fondation Bristol Myers Squibb pour la Recherche en Immuno-Oncologie (BMS 2104009NA); French National Cancer Institute (INCa), Fondation ARC, and Ligue Nationale contre le Cancer (PAIR TUMC21-001, INCa-16280); Entreprises contre le Cancer Paris-Île-de-France (GEFLUC R20202DD); and TRANSCAN3 (C20/BM/14646004/GLASSLUX, INTER/TRANSCAN22/17612718/PLASTIG). A.L. was supported by a Fondation pour la Recherche Médicale (FRM) fellowship, Q.R. by a fellowship from the French Ministry of Education and Research, and Y.H. by a fellowship from La Ligue Nationale contre le Cancer.

## Methods

### Human samples and CD11B-based sorting

Brain tissues were collected following written informed consent under protocols approved by the Institutional Ethical Committee Board in accordance with the Declaration of Helsinki. Samples for analyses were selected from the Pitié-Salpêtrière tumor bank Onconeurotek based on clinical information and validation by expert neuropathologists (K.M., S.T., F.B) of both histological features and molecular diagnosis as previously described^81^. Cryopreservation was performed as follows: tumors taken from the neurosurgery room were immediately transported on ice in HBSS (Gibco), cut into 2-5 mm diameter pieces, and submerged in cryotubes containing 1 mL of DMEM/F-12 (Gibco), 20% FBS, and 10% DMSO (Sigma-Aldrich). The cryotubes were placed at −80°C in alcohol-free freezing containers (Corning® CoolCell®). Cryopreserved samples were stored in liquid nitrogen until use. On the day of analysis, the tumor pieces were thawed and rinsed in DMEM/F-12. Next, the tissue was enzymatically and mechanically digested for 5-10 minutes at 37°C in HBSS-papain-based lysis buffer (Worthington) containing DNAse (0.01%, Worthington) and L-cysteine (124 μg/mL, Sigma). Enzymatic digestion was stopped with ovomucoid (700 μg/mL; Worthington). The homogenates were sequentially filtered using 100 μm, 70 μm, and 30 µm SmartStrainers (Miltenyi) to remove residual clumps. Cell pellets were resuspended in an appropriate volume of cold MACS debris removal solution according to the manufacturer’s instructions (Miltenyi). The suspension was gently mixed by pipetting, and ice-cold DPBS was added above the cell suspension to create a transparent gradient. The suspension was centrifuged and resuspended in cold MACS Buffer (0,5% BSA and 2 mM EDTA in PBS). After centrifugation, the supernatant was completely removed and red blood cell lysis was performed using ACK buffer (Gibco) for 5 min at room temperature. Next, the cells were centrifuged at 4°C and pellets were washed with cold DPBS. After, the cells were resuspended in MACS buffer and counted, and the cell pellets were labeled with an appropriate volume of CD11B MicroBeads (Miltenyi) in MACS buffer and incubated for 15 min at 4°C. Subsequently, the cells were resuspended in cold MACS buffer and centrifuged at 300 g for 10 min at 4°C. The supernatant was aspirated, and the cells were resuspended in cold MACS buffer. The LS MACS Separation Columns were placed in the magnetic field of a suitable MACS Separator, and the cell suspension was applied to the columns. The CD11B^+^ and CD11B^−^ fractions were collected separately on ice. The dry pellets were stored at −80°C until nucleic acid extraction.

### Nucleic acid extraction

RNA and DNA from CD11B^+^ and CD11B^−^ samples were co-eluted using the AllPrep DNA/RNA Micro Kit (Qiagen, 80284) or AllPrep DNA/RNA Mini Kit (Qiagen, 80204), following the manufacturer’s instructions. In some cases, nucleic acids were purified separately using the Maxwell RSC simplyRNA Cells Kit (Promega, AS 1390) and Maxwell RSC Blood DNA Kit (Promega, AS 1400) following the manufacturer’s instructions. The yield and quality of total RNA were assessed using a Tapestation 2200 (Agilent) instrument.

### Digital droplet PCR

IDH1 R132H, TERT C228T, and TERT C250T mutations were analyzed in the CD11B^+^ and CD11B^−^ fractions from human gliomas using ddPCR. Droplet generation and partitioning were performed using the Bio-Rad system QX200. Primer sets, probes, and ddPCR Supermix for Probes (No dUTP) were used to quantify the absolute copy number in 3 ng of input DNA following the manufacturer’s instructions. All the primers and probes were obtained from Bio-Rad Laboratories. Sequence information is available at www.bio-rad.com using the ID numbers: IDH1 p.R132H Hsa, Human (Ref: 10031246 UniqueAssayID: dHsaCP2000055), IDH1 WT for p.R132H Hsa, Human (Ref: 10031249 UniqueAssayID: dHsaCP2000056), TERT C228T_113 Hsa, Human (Ref: 12009308 UniqueAssayID: dHsaEXD72405942), TERT C250T_113 Hsa, Human (Ref: 12003908 UniqueAssayID: dHsaEXD46675715). The data were analyzed using Poisson statistics to determine the target DNA template concentration in the original sample. Positive controls consisted of tumor DNA from the CD11B^−^ fractions, and negative controls contained water instead of DNA.

### DNA methylation profiling

Genomic DNA was quantified using the Quant-iT dsDNA Broad Range Assay (Thermo Fisher Scientific, Waltham, MA, USA) on a Tecan SPARK microplate reader (TECAN, Switzerland). Total DNA (500 ng) from each sample was sodium bisulfite-converted using the Zymo EZ-96 DNA Methylation kit following the manufacturer’s recommendations for the Infinium assay (Zymo Research, catalog number: D5004). After conversion, the DNA concentration of each sample was adjusted to 50 ng/µL with M-elution Buffer or concentrated using speed vacuum. In total, 300 ng of converted DNA for each sample was used as a template on the Infinium MethylationEPIC BeadChip following the manufacturer’s recommendations. Briefly, bisulfite-converted DNA was whole-genome amplified, fragmented, and hybridized to the array. After hybridization, the unhybridized and non-specifically hybridized DNA was washed away and the captured product was extended with fluorescent labels coupled to nucleotides. Finally, the arrays were scanned using a high-resolution Illumina scanner (iScan), which acquires light images emitted from the fluorophores. The intensities were measured and methylation signals were extracted and recorded as raw data (IDAT). These assays were performed at the P3S platform (Sorbonne University).

DNA methylation data were analyzed using the RnBeads package. Briefly, after the removal of bad quality samples, data were aligned to the hg19 assembly using the beta Mixture Quantile dilation (BMIQ) method for normalization and adjusted for sex as a covariate. Sex chromosomes were included, and sites with a standard deviation below 0.01 across all samples were filtered out. Together with default annotations, we included custom enhancer annotations corresponding to entire enhancer regions defined by the FANTOM5 enhancer-robust promoter association dataset. Briefly, enhancer–promoter pairs located within 500 kb were identified by correlating enhancer and promoter activity across cell types and organs, and associations passing an adjusted Pearson correlation threshold of FDR < 1 x 10^−5^ were retained. These associations were computed between FANTOM5 enhancers and all FANTOM robust promoters and were used as putative enhancer-gene links. Methylation-based deconvolution was performed using the Glioma Immune Microenvironment Composition Calculator (GIMiCC) on MethylationEPIC array data^82^. GIMiCC estimates glioma microenvironment composition from DNA methylation profiles using reference methylomes from isolated CNS and immune cell populations and a hierarchical deconvolution framework, as previously described. The non-tumor microglial reference used in the original GIMiCC framework included CD11B⁺ fractions that were also analyzed in the present study^33^. Estimated proportions of microglia, monocyte/macrophage-lineage cells, and other minor populations were extracted from CD11B⁺ myeloid fractions and compared according to tumor IDH status. Epigenetic clock analysis was conducted using “epiTOC2” (Epigenetic Timer Of Cancer) which calculates the mitotic age of cells based on methylation levels of 163 CpG sites from the EPIC arrays^83^.

### RNA sequencing

Library preparations were performed with Kapa mRNA Hyper prep (Roche) following the manufacturer’s instructions and sequenced (paired end) with the Illumina Novaseq 6000 Sequencing system with 200 cycles cartridge to obtain 2 × 60 million reads 100bases/RNA. Sequencing was performed at the iGenseq platform at the ICM – Paris Brain Institute. For the *in vitro* experiments, we combined Illumina Stranded Total RNA Prep and Ribo-Zero Plus. The quality of the raw data was evaluated using Fast QC. Sequences were trimmed or removed using Fastp software to retain only high-quality paired reads. Star v2.5.3a was used to align reads on the GRCh38 reference genome using default parameters, except for the maximum number of multiple alignments allowed for a read set to 1. Transcript quantification was performed using the rsem 1.2.28. Differential expression analysis was conducted using the quasi-likelihood F-test in the edgeR package. Normalized data were computed using the cpm function. Multiple hypothesis testing was corrected using the False Discovery Rate Benjamini-Hochberg method. Samples from all *in vitro* experiments were analyzed in a paired manner for each individual.

### Targeted proteomics

Cell pellets from freshly isolated CD11B^+^ myeloid fractions were lysed in 50-200 µL RIPA lysis buffer (Millipore, #20-188). Protein abundance of 1,034 inflammation- and immune-related targets was profiled using the Olink Reveal platform (proximity extension assay, PEA, with next-generation sequencing readout). Cell lysates (but not cell culture supernatants) were diluted to 0.5 mg/mL total protein and 4 µL per sample was analyzed on randomized 96-well plates together with Olink internal synthetic controls for incubation, extension, and amplification steps. Sequencing read counts were processed using Olink’s standard normalization pipeline to generate normalized protein expression (NPX) values on a log2 scale. Proteins were retained for differential expression analyses only if NPX values were above the assay limit of detection (LOD) in the relevant comparison groups. Group differences are reported as ΔNPX (treated-control), corresponding to log2 fold change. Proteins were considered differentially expressed at |ΔNPX| > 0.5 and FDR < 0.05.

### LC-MS/MS quantification of genomic 5mC and 5hmC

Genomic DNA (gDNA) was isolated as indicated and concentrated by ethanol precipitation. Samples were then digested at 37°C for 12 h in 25 µL reactions containing 1-4 µg of gDNA, 7.5 U of DNA Degradase Plus (Zymo Research), reaction buffer, and isotope-labeled internal standards [700 pmol 2’-deoxycytidine-15N3 (Cambridge Isotope Laboratories, Inc.); 1.75 pmol 5-methyl-2’-deoxy cytidine-d3 (Toronto Research Chemicals), 1 pmol 5-hydroxymethyl-2’-deoxycytidine-d3 (Toronto Research Chemicals)] for quantification. Prior to LC/MS analysis, the samples were diluted 1:1 in mobile phase component A (see below). The LC-MS system consisted of a Shimadzu Nexera UPLC system in line with a SCIEX 6500 QTrap mass spectrometer equipped with a TurboV Ion spray source and operated in positive ion mode. The analytes were chromatographed on a gradient elution system in which mobile phase A was water with 10 mM ammonium formate and 0.05% trifluoroacetic acid (TFA) (Sigma Aldrich), and mobile phase B was 7:1 methanol:acetonitrile (Thermo Fisher Scientific) with 10 mM ammonium formate and 0.05% TFA. The column was a Phenomenex Curosil-PFP column (10 x 0.2 cm 1.7 um) (Phenomenex Inc.) held at 42°C. The flow rate was 300 µL/min and the gradient started at 1.0% B (held for 30 s) and was then increased to 19% over 5 min. All analytes were detected using selected reaction monitoring (SRM). Analyst software (version 1.6.2) was used to acquire and process data. The analytes were quantified using stable isotope dilutions against stably labeled internal standards. Data represent the percentage of each analyte relative to the total cytosine pool (dC+5mC+5hmC) in the same sample.

### Sequence motif enrichment analysis

Motif enrichment was performed using Hypergeometric Optimization of Motif EnRichment (HOMER) v4.11^84^, enabling masking repeats, setting the parameter size to “given” and searching for motifs with a length within the range of 7-14 bp. We used HOCOMOCO v12 CORE as the reference dataset (matrix with a threshold level corresponding to a p-value of 0.001)^85^.

### Single-sample Gene Set Enrichment Analysis (ssGSEA)

Single-sample Gene Set Enrichment Analysis (ssGSEA) scores were computed for predefined inflammatory and suppressive transcriptional programs (“CXCR4 Inflam.”, “IL1B Inflam.”, “Scav.Suppres.”, “C1Q Suppres.”, and “IFN Response”)^31^, across all CD11B^+^ myeloid fractions from the *ex vivo* study. Differences in program activity between IDH-mutant, IDH-wt, and non-tumor groups were assessed using pairwise Wilcoxon rank-sum tests implemented in the rstatix R package. P-values were adjusted for multiple testing using the Benjamini–Hochberg (BH) method. Enrichment scores were visualized using boxplots generated with ggplot2, and adjusted p-values were displayed using ggpubr function stat_pvalue_manual.

### Regulon analysis

To infer transcription factor activity, we used the CollecTRI database^45^, which contains regulons comprising signed TF-target gene interactions, to infer TF activity. To estimate TF activity in hypermethylated enhancers, we considered genes with associated overexpression or downregulation (falling in quadrants I and IV, respectively; Fig. 2a). These genes were then interrogated among the targets of each TF in the database. TFs with at least five targets that were regulated in a positive or negative manner were retained. We then computed TF activity using the Normalized Enrichment Score (NES) of the Mann-Whitney-Wilcoxon gene set test (MWW-GST)^46^. We applied MWW-GST to each sample’s normalized data and retained TFs with an FDR < 0.05, for the NES in at least 18 samples and an FDR < 0.05, in the comparison of the activity between groups (Wilcoxon’s test). Heatmap depicting TF activity at hypermethylated enhancers and promoters was generated using the z-score of activity.

### Primary human microglial cultures

Primary microglial cultures were established using cavitron ultrasonic surgical aspirates (CUSA) obtained from patients undergoing surgery from brain tumors or drug-resistant epilepsy. Brain tissues were collected following written informed consent under protocols approved by the Institutional Ethical Committee Board. Tissue source, IDH status, downstreams applications performed using each culture, and the pathological, molecular and demographic characteristics of the cohort are summarized in Supplementary Table S1. Independent biological replicates corresponded to cultures derived from distinct donors. Upon receipt (averaging 2 h), the tissues were promptly processed according to the protocol specified for human tumor samples. After enzymatic digestion and MACS purification, CD11B^+^ cells were counted and directly seeded in non-coated 12-well plates at a density of 750-1000 K cells/well in DMEM/F12, GlutaMAX medium (Gibco Ref# 10565018) supplemented with N-acetyl-cysteine and insulin [5 µg/mL], apo-Transferrin [100 µg/mL], sodium selenite [100 ng/mL], human TGFβ2 [2 ng/mL], CSF-1 [20 ng/mL], and ovine wool cholesterol [1.5 µg/mL]^55^. In addition, cultures were supplemented with ascorbic acid [75 µM]. All reagents and growth factors were purchased from Sigma and Preprotech, respectively. Non-cell-permeable D-2HG acid disodium salt (Sigma-Aldrich) was used at a final concentration of 5 mM for all experiments, with treatments starting after 48 h of culture. The concentration was selected because it approximates concentrations previously reported in IDH-mutant glioma microenvironments^5,6^. Half of the culture medium was renewed every 2-3 days. For the 6 h LPS challenge experiments, 1% FBS was added to all conditions.

### IDH-wt and IDH-mutant cell lines

Patient-derived IDH-mutant glioma cells were kindly provided by Artee Luchman and Sam Weiss (University of Calgary) and Keith Ligon (Dana-Farber Cancer Institute), whereas IDH-wt glioma cells were established by the GlioTEx team (Glioblastoma and Experimental Therapeutics) at the ICM-Paris Brain Institute. Cells were maintained as neurospheres in DMEM/F12 supplemented with B27, EGF (20 ng/mL), FGF (20 ng/mL), and 1% penicillin/streptomycin at 37 °C in a humidified atmosphere containing 5% CO₂. Conditional Idh1 knock-in neural stem cells (NSCs) were generated from postnatal (P0-P1) mice carrying the conditional Idh1 R132H allele crossed with *Rosa26^LSL-YFP^*. NSCs were cultured under standard neurosphere conditions in serum-free DMEM/F-12 GlutaMAX™ medium supplemented with B27, N2, 0.6% w/v D-glucose, 5mM HEPES, and 20μg/ mL insulin. Cre-mediated recombination was induced using a Cre-expressing adenovirus (VectorBuilder), leading to expression of Idh1 R132H and YFP. Efficient recombination was verified by YFP fluorescence. Cells were maintained for 3–6 passages and used as positive (IDH-mutant) and negative (IDH-wt) controls in fluorometric assays measuring intracellular D-2HG levels. Human immortalized microglial cells (HMC3; ATCC® CRL-3304™) were maintained in low-glucose DMEM containing L-glutamine supplemented with 10% FBS. Cells were treated with cell-permeable octyl-D-2HG (350 µM) or vehicle control (0.3% DMSO) for up to 30 days. For 6 h LPS challenge experiments, cells were switched to DMEM supplemented with 0.2 mM glutamine, and the same additives used for primary microglial cultures including 1% FBS which was added to all conditions.

### Immunofluorescence staining

Cells were seeded in 12 multiple wells at a density of 750 K cells/well and maintained as described for 3, 10, and 17 days. The cells were washed with PBS (Gibco) and fixed with 4% paraformaldehyde (Electron Microscopy Sciences) in PBS for 10 min at RT. The cells were washed with PBS and permeabilized with 0,1% Triton X-100 in 1x PBS for 5 min at RT. Cells were quickly washed with PBS and blocked for non-specific binding sites with Human Fc Block (#564219, BD Pharmingen) for 10 min at RT, and subsequently with 5% donkey in PBS for 30-60 ²min at RT. Incubation with primary antibodies was performed overnight at 4°C using anti-Iba1 (#19-19741, Wako; diluted 1/800), Tmem119 (#A16075D, BioLegend; diluted 1/100), or Ki-67 (#AB16667-1001, Abcam; diluted 1/250) in a humidified and light-protected chamber. The cells were washed with PBS and incubated with secondary antibodies (anti-rabbit #A21206 or anti-mouse #A21203; diluted 1/1000) in a blocking solution for 30 min at RT and temperature in the dark. Subsequently, the cells were washed with PBS, mounted with Fluoroshield-DAPI solution (Sigma), and observed under an Apotome inverted microscope (ZEISS Apotome 3) and a confocal microscope (A1R HD25 Nikon Inverted). Negative controls (background fluorescence) were obtained by omitting the primary antibodies. The percentages of TMEM119^+^ and IBA1^+^ cells were determined in primary cultures grown in uncoated wells by counting three to five fields covering 0.22% of the well surface at different time points over a two-week period. Fields were randomly selected in the area where the cell density, as assessed by DAPI staining, was adequate.

### Quantification of D-2HG levels

Cultured cells were pelleted by centrifugation to remove the media, followed by two washes with ice-cold DPBS to ensure the complete removal of all D-2HG-containing media. Microglial cells were collected by trypsinization (0.25% for 5 min), whereas glioma cells and neurospheres were collected using Accutase. The corresponding culture medium, supplemented with 10% FBS, was added for inactivation. After centrifugation (300 g at 4°C for 5 min), the resulting pellet was washed with cold DPBS. Cell pellets were then preserved at −80°C until use. The concentration of D-2HG in the cell lysates was determined using the D-2-Hydroxyglutarate (D-2HG) Assay Kit (Ref MAK320, Sigma) and LC/MS for validation. For the fluorometric assay, cellular extracts were prepared by adding 75 µL of CelLytic MT Cell Lysis Reagent (Sigma) with a Protease Inhibitor Cocktail (Sigma). An aliquot was used for protein quantification using the Pierce™ BCA Protein Assay Kit (Thermo Fisher Scientific). The collected supernatants were deproteinized by perchloric acid precipitation and neutralized with KOH. A standard curve was prepared using serial dilutions of D-2HG. An equivalent of 5000 pmol was added to the cell extract and used as a spike-in internal control. The plates were then incubated at 37°C for 30–60 min. This assay is based on the oxidation of D-2HG to α-ketoglutarate (αKG) by the enzyme (D)-2-hydroxyglutarate dehydrogenase (HGDH) coupled with the reduction of NAD+ to NADH^86^. The amount of NADH formed was then quantitated by diaphorase-mediated reduction of resazurin to the fluorescent dye resorufin (λex = 540 nm, λem = 590 nm) using a Spectramax i3x microplate reader. The concentration of D-2HG in each sample was estimated as pmol/µg protein. For quantification of 2HG using LC-MS, dry cell pellets were lysed in water, and the solution was divided in half for 2-HG measurement and protein quantification. NaCl was added and the samples were acidified using HCl. Liquid-liquid extraction of organic acids with ethyl acetate was performed. Three extractions were performed and the organic phases were pooled and dried under a nitrogen stream at 30°C. The samples were then derived using a standard silylation protocol, N, O-bis [trimethylsilyl] trifluoroacetamide, and 1% trimethylchlorosilane (TMCS) under anhydrous conditions using pyridine. Chromatographic separation was performed using a TR-5MS (30m x 0.25 mm x 0.25 mm) column from Thermo Scientific (Waltham, Massachusets, USA). The spectral data were acquired using XCalibur software (Thermo Electron Corporation, Austin, TX, USA). The samples were placed for 30 min at 80°C and then injected into the GC system. Quantification was performed using the internal standard 2-hydroxyglutaric acid-D3 from Cambridge Isotope Laboratories (Tewksbury, Massachusetts, USA).

### TET enzymatic assay

Nuclei were isolated using a nuclear extraction kit (Cat. Ab113474, Abcam, Cambridge, MA, USA). Cells were collected by trypsinization (0.25% for 5 min) and counted. The cells were then resuspended in an appropriate volume of pre-extraction buffer containing dithiothreitol and protease inhibitor cocktail and incubated on ice for 2 min. After centrifugation at 12,000 rpm for 3 min, the cytoplasmic extract (supernatant) was carefully removed, leaving a nuclear pellet. Two volumes of extraction buffer containing dithiothreitol and protease inhibitor cocktail were added to the nuclear pellet. The extract was incubated on ice for 15 min and vortexed every 3 min for 5 s. After incubation, the suspension was centrifuged for 10 min at 14,000 rpm and 4°C, and the resulting supernatant was transferred to a new microcentrifuge vial. Protein quantification from the nuclear extract was performed using a Pierce™ BCA Protein Assay Kit (Thermo Fisher Scientific) following the manufacturer’s instructions. Technical triplicates for this assay were performed using 12 µg of protein from nuclear extracts of microglial cells (untreated and treated with D-2HG), as well as from external controls (neurospheres obtained from the cKI Idh R132H mouse model). To determine TET enzymatic activity, we used the TET Hydroxylase Activity Kit from Abcam (ab156913). In this fluorometric assay, a methylated substrate was stably coated onto microplate wells. Active TET enzymes bind to the substrate and convert the methylated substrate into hydroxymethylated products. The hydroxymethylated products generated by the TET activity can be detected using a specific antibody. The ratio or amount of hydroxymethylated products, which is proportional to the enzyme activity, was measured fluorometrically using a fluorescent microplate reader with excitation at 530 nm and emission at 590 nm. The TET enzyme activity was proportional to the measured relative fluorescent units.

### Reduced Representation Enzymatic Methyl sequencing (RREM-seq)

Library preparation was performed using 100 ng of genomic DNA spiked with 0.95 ng of unmethylated Lambda DNA and 0.05 ng and CpG methylated pUC19. The mixture was digested with 400 units of MspI (New England Biolabs) in NEB Ultra II buffer (New England Biolabs) in a final volume of 57 µL for 2 h at 37°C. After enzymatic DNA digestion, the fragments were repaired and A-tailed using 15 units of Klenow 3’-5’ exo-(New England Biolabs). After End-repair and A-tailing, Illumina universal methylated adapters were ligated to DNA fragments using the NEB Ultra II ligation module, following the manufacturer’s recommendations. After ligation, DNA fragments were purified and size-selected using two-step SPRI bead purification to select fragments from 40bp to 400bp. The clean-up ligated DNA was eluted with 10 mM Tris at pH 8. The purified product was then converted using the enzymatic methyl-seq module (New England Biolabs), following the manufacturer’s recommendations. Briefly, 5mC was converted to 5hmC by TET2, and 5hmC was specifically protected from APOBEC deamination by T4-BGT glycosylation. After 5hmC/5mC glycosylation, the libraries were denatured for 10 min at 85°C in 20% formamide. The unmodified cytosines were then deaminated during the APOBEC reaction. The converted product was amplified by PCR using indexing primers and polymerase KAPA HiFi Hotstart U+ Ready Mix 2X in a final volume of 50 µL for 13 cycles. PCR products were purified at a 1.2 × SPRI ratio and then controlled by capillary electrophoresis on a Fragment Analyzer. After quantification by qPCR, libraries were sequenced on Illumina Novaseq 6000 paired-end 100b reads.

### Reduced Representation Enzymatic hydroxyMethyl sequencing (RREhM-seq)

We used the RREhM-seq protocol described by Sun et al^58^ with slight modifications. Library preparation was performed using an input of 200 ng with a spike of 1.9 ng of unmethylated Lambda DNA and CpG methylated pUC19 (0.1 ng). Following MspI digestion and reparation with Klenow 3’-5’ exo- and pyrrolo-dC, Illumina adapters were used instead of 5mC adapters during the ligation step. For the TET2 step during conversion, the TET2 enzyme and Fe2+ were replaced with H_2_O to avoid 5mC protection. The product was deaminated, amplified, qualified, and sequenced under the same conditions as those used in the RREM-seq libraries. RREM-seq and RREhM-seq were performed at Integragen SA (Evry, France).

### RREM-seq & RREhM-seq analysis

The FastQ files were aligned using BSseeker2 (https://github.com/BSSeeker/BSseeker2) on the hg19 genome using bowtie2. The hg19 index was built with the rrbs option, and fragment size between 40bp and 400 bp were selected. The adapter sequence and CCGG motifs were removed during alignment. After alignment, methylation calling was performed to obtain the methylation level and cover each cytosine position in the library. Options for removing overlaps and low-quality sequences were also enabled. For RREM-seq libraries, methylation signals correspond to 5mC and 5hmC, whereas for RREhM-seq libraries these only concern to 5hmC^84^. Total 5mC (5mC + 5hmC) and 5hmC levels were calculated as the percentage of modified reads among all reads covering a given cytosine. The 5mC-only level was calculated by subtracting the 5hmC level from the total 5mC level. CpG sites were analyzed with the methylKit package^87^ using a minimum coverage of 10 reads per base. For the defined window analysis, TFs motifs were retrieved from HOCOMOCO v12 CORE and searched inside entire enhancer regions using the Biostrings package^88^ with a maximum mismatch of one base. The region of interest was defined as a 600-bp window centered on the first base of each motif. Motif analyses were restricted to enhancers previously identified as hypermethylated in CD11B⁺ fractions from IDH-mutant gliomas relative to non-tumor brain tissues. Microglial samples used for 5mC and 5hmC analyses at 14 days were obtained from surgical aspirates of non-tumor brain tissues from three patients who underwent surgery for epilepsy. One technical outlier from the 5hmC analysis, identified by aberrantly low sequencing coverage across enhancer regions, was excluded. The 5mC-only and 5hmC levels were averaged across patients for each condition using only the common enhancer regions or defined windows shared among patients.

### Orthotopic glioma xenograft and chronic D-2HG infusion

Six-week-old male athymic nude mice were acclimatized for at least 14 days prior to surgery. All animal procedures were performed at the Institute of Psychiatry and Neurosciences of Paris (IPNP) in accordance with approved APAFIS protocol no. 2021101316589067 (“Effets du D-2 HydroxyGlutarate et des mutations de l’enzyme Isocitrate DesHydrogénase sur la croissance des gliomes : xenogreffe intracérébrale de lignées cellulaires de glioblastome”). Patient-derived glioblastoma cells (GLTX-20; GlioTEx) expressing GFP were used for orthotopic xenografting. Mice received a stereotaxic injection of 3 × 10⁵ GLTX-20-GFP cells suspended in 4 µL DMEM/GlutaMAX medium into the right striatum (AP +0.3 mm, ML +1.5 mm, DV −1.25 mm). All surgical procedures were performed under isoflurane anesthesia (3% for induction and 1.5% in 1 L/min O₂ for maintenance). Animals were maintained on a heating pad with rectal temperature monitoring, and ophthalmic gel was applied to prevent corneal dehydration. Pre-operative analgesia was provided with buprenorphine (0.1 mg/kg, subcutaneous), and local anesthesia was achieved by application of diluted lidocaine to the skull. To compensate for fluid loss, 250 µL sterile 0.9% saline was administered subcutaneously. Seven days after tumor-cell injection, mice underwent a second stereotaxic procedure for implantation of an iPRECIO osmotic pump enabling continuous intracerebral delivery of D-2HG or saline control. The cannula was inserted through the original burr hole and secured to the skull with surgical adhesive. Pumps were filled with 25 mM D-2HG in sterile saline and programmed to deliver the solution at 0.1 µL/h for 14 days. Post-operative analgesia was provided with meloxicam (Metacam; 20 mg/kg) immediately after surgery and again 24 hours later. Animals were monitored daily throughout the study for body weight, general condition, and signs of pain or distress. No animals required euthanasia because of postoperative complications. Fourteen days after pump implantation, mice were euthanized and tumor-bearing brains were harvested. CD11B⁺ myeloid cells were isolated from dissociated brain tissue as described above. Genomic DNA from purified CD11B⁺ cells was subjected to RREhM-seq library preparation and analysis following the procedures described above.

### RT-qPCR

RNA from cultured microglia was isolated using a Maxwell RSC simplyRNA Cells Kit (Promega). A total of 300 ng of RNA was reverse-transcribed into complementary DNA (cDNA) using the Maxima First Strand cDNA Synthesis Kit (Thermo Fisher Scientific). RT-qPCR assays were performed in technical replicates on a Light Cycler 480 instrument using SYBR Green I Master Mix (Roche). Primer sequences were as follows:

*TNFA*, forward: 5’-GAGCCAGCTCCCTCTATTTA-3’; reverse: 5’-GGGAACAGCCTATTGTTCAG-3’; *IL6*, forward: 5’-CCTTCCAAAGATGGCTGAAA-3’; reverse: 5’-TGGCTTGTTCCTCACTACT-3’; *PPIA*, forward: 5’-ATGCTGGACCCAACACAAAT-3’; reverse: 5’-TCTTTCACTTTGCCAAACACC-3’; *ACTB*, forward: 5’-CCAACCGCGAGAAGATGA-3’; reverse: 5’-CCAGAGGCGTACAGGGATAG-3’. Primer linearity, amplification efficiency, and specificity were verified for all assays. Relative cytokine expression levels were normalized to the mean expression of the housekeeping genes *PPIA* and *ACTB* and expressed as fold change.

### Mitochondrial respiration measurement

Real-time measurement of the oxygen consumption rate (OCR) was performed using an XFe extracellular flux analyzer (Seahorse Bioscience, Billerica, MA, USA), following the manufacturer’s instructions. Primary isolated microglia were plated at a density of 1,000,000 cells/well in XFe24 cell culture microplates pre-coated with poly-L-lysine (Sigma-Aldrich). Cells were maintained in 200 μl DMEM/F-12 medium, supplemented as previously indicated, and treated with 5 mM D-2HG for 12 days. Half of the culture medium was renewed every 2-3 days. On the day of the experiment, cells were stimulated with LPS (1µg/mL) in medium supplemented with FBS 1% for 6 h. FBS 1% was also added to the other experimental conditions. Next, the medium was replaced by Seahorse XF base medium (Agilent, 102353-100) supplemented with 17.5 mM D-Glucose (Dextrose), 0.5 mM sodium pyruvate and 2.5 mM L-Glutamine and incubated for 1 h in CO2 free incubator at 37°C. The utility plate was hydrated with an XF calibrant solution (pH 7.4) (Agilent, 100840-000; 1 mL/well) and incubated overnight (37°C, CO2-free). The next day, the utility plate with the cartridge containing the injector ports and sensors was run on Seahorse for calibration. The assay medium (Seahorse XF DMEM assay medium, pH 7.4) was prepared immediately before the assay. OCR was measured under basal conditions and after sequential injections of the following inhibitors: oligomycin (1 μM) to stop ATP synthase, FCCP (1 μM) to dissipate the proton gradient, and Rotenone/Antimycin A (0.5 μM each to block the electron transport chain (ETC). All the following measurements were carried out with a 2 min mix, 2 min delay, and 3 min measure. Three baseline OCR measurements were recorded, and mitochondrial metabolism was assessed by injection of oligomycin, FCCP, and a combination of rotenone and antimycin A. The following parameters were deduced: basal respiration (OCR values used to provide ATP under baseline conditions), ATP-linked respiration (following oligomycin injection, a reduction in OCR values represents the part of basal respiration used to produce ATP), and maximal respiration (maximal OCR values following FCCP injection). Basal respiration was calculated by subtracting the last rate measured before oligomycin injection from the non-mitochondrial respiration rate (NMRT), which was determined as the minimum rate measured after rotenone/antimycin A injection. Maximal respiration was calculated by subtracting the maximum measurement after the FCCP injection and NMRT. ATP-linked respiration was calculated by subtracting the last rate measured before oligomycin injection from the minimum rate measured after oligomycin injection. OCR data were normalized to the number of viable cells estimated at the end of the experiment. To do this, the cells were fixed with paraformaldehyde 4%, followed by washing with PBS and labeling with DAPI [10 µg/mL]. Nuclei were automatically counted on the CellInsight NXT platform.

### Single-nucleus RNA sequencing and data processing

Information for the three patients analyzed in this study are presented in Supplementary Table S1. Matched pre- and post-treatment samples from Patient 02 were processed using the Parse Biosciences Evercode Whole Transcriptome platform (v3 chemistry). Nuclei were isolated by detergent-based lysis, fixed using Parse Evercode Cell Fixation v3 (Parse Bioscience, Seattle, WA, USA), and approximately 8,000 nuclei per sample were loaded into each split-pool barcoding reaction. Barcoding, reverse transcription, library preparation, and PCR amplification were performed according to the manufacturer’s instructions. Libraries were pooled and sequenced on an Illumina NovaSeq instrument at the iGenseq platform (ICM, Paris Brain Institute), yielding a sequencing depth of approximately 43,000 reads per nucleus. Raw FASTQ files were processed using Split-pipe v1.6.1 in “all” mode with v3 chemistry and the WT MEGA kit. A custom reference index was generated using the “mkref” function with Ensembl release 109 annotations. Split-pipe performed adapter trimming, alignment, UMI and barcode error correction, and PCR duplicate removal. Individual sublibraries were subsequently merged using the “combine” function to generate experiment-wide count matrices. Matched pre- and post-treatment samples from Patient 01 (BWH445) and Patient 03 (PARIS) were generated using the 10x Genomics platform in a previous study^61^. Downstream analyses were performed using Seurat v5.3. For the Parse dataset, nuclei with fewer than 500 or more than 10,000 detected genes (nFeature_RNA), fewer than 500 or more than 30,000 unique molecular identifiers (UMIs; nCount_RNA), or greater than 5% mitochondrial reads were excluded. For the 10x genomics dataset, cells were retained if they contained 500–6,500 detected genes (nFeature_RNA), 1,000–35,000 UMIs (nCount_RNA), and ≤15% mitochondrial reads. Doublets were identified and removed from each sample individually using scDblFinder. Data were normalized using SCTransform while regressing out mitochondrial content (percent.mt). Principal component analysis (PCA) was performed using 3,000 integration features, and batch effects were corrected using Harmony, grouping by sample identity. To avoid confounding effects related to sequencing chemistry or platform, Parse and 10x Genomics datasets were analyzed separately. To enable consistent identification of microglial populations across patients, the processed Parse and 10x genomics datasets were subsequently merged after independent normalization and batch correction. All high-quality, doublet-removed, Harmony-integrated samples were combined using Seurat’s merge function for unified identification and analysis of microglial populations.

### snRNA-seq analysis of microglial cells

The combined datasets were clustered based on the Harmony-corrected embeddings using a shared nearest-neighbor (SNN) graph, followed by UMAP visualization. Microglia were computationally identified using UCell enrichment scores of a microglial gene signature comprising TMEM119, P2RY12, SALL1, GPR34, P2RY13, SLC2A5, and TAL1. The cluster showing the highest enrichment score for this signature was annotated as microglia and extracted for downstream analyses. For Patient P02, microglia nuclei were log-normalized, 2,000 highly variable genes were selected, and the data were scaled prior to PCA. The first 15 principal components were used to construct the neighbor graph and compute UMAP embeddings. Differential gene expression analysis between pre- and post-treatment microglia was performed using Seurat’s FindMarkers function (Wilcoxon rank-sum test; min.pct = 0.05, logfc.threshold = 0), with Bonferroni correction. Genes with log2 fold change > 1 and adjusted P < 0.05 were considered significant. An analogous workflow was applied to microglia from Patients 01 and 03, with dataset-specific dimensionality reduction and clustering parameters as appropriate. All tested genes for each patient are shown in Supplementary Table S8.

### Statistical analysis and data visualization

Sample sizes and statistical tests are indicated in the corresponding figure legends. All statistical tests were two-sided unless otherwise specified. P-values were adjusted for multiple testing using the Benjamini–Hochberg (BH) procedure, with a significance threshold of α = 0.05. Statistical analyses were performed in R version 4.4.1 (R Foundation for Statistical Computing, Vienna, Austria). Normality was assessed using the Shapiro–Wilk test. Data visualization was performed using BioRender and the R packages ggplot2, ggsignif, ComplexHeatmap, clusterProfiler, and eulerr. Gene regulatory network was visualized with Cytoscape. The Hallmark gene set collection (v2023.1.Hs) used for enrichment analyses was obtained from the Molecular Signatures Database (MSigDB)^89^.

### Data and code availability

The datasets generated during the current study are currently being deposited in a public repository. Accession numbers will be provided in the final version of the manuscript. All newly generated data and the custom code used for data processing and analysis will be made publicly available upon publication, or earlier if the deposition process is completed before acceptance.

## Supplementary Figures

**Supplementary Fig. S1.**
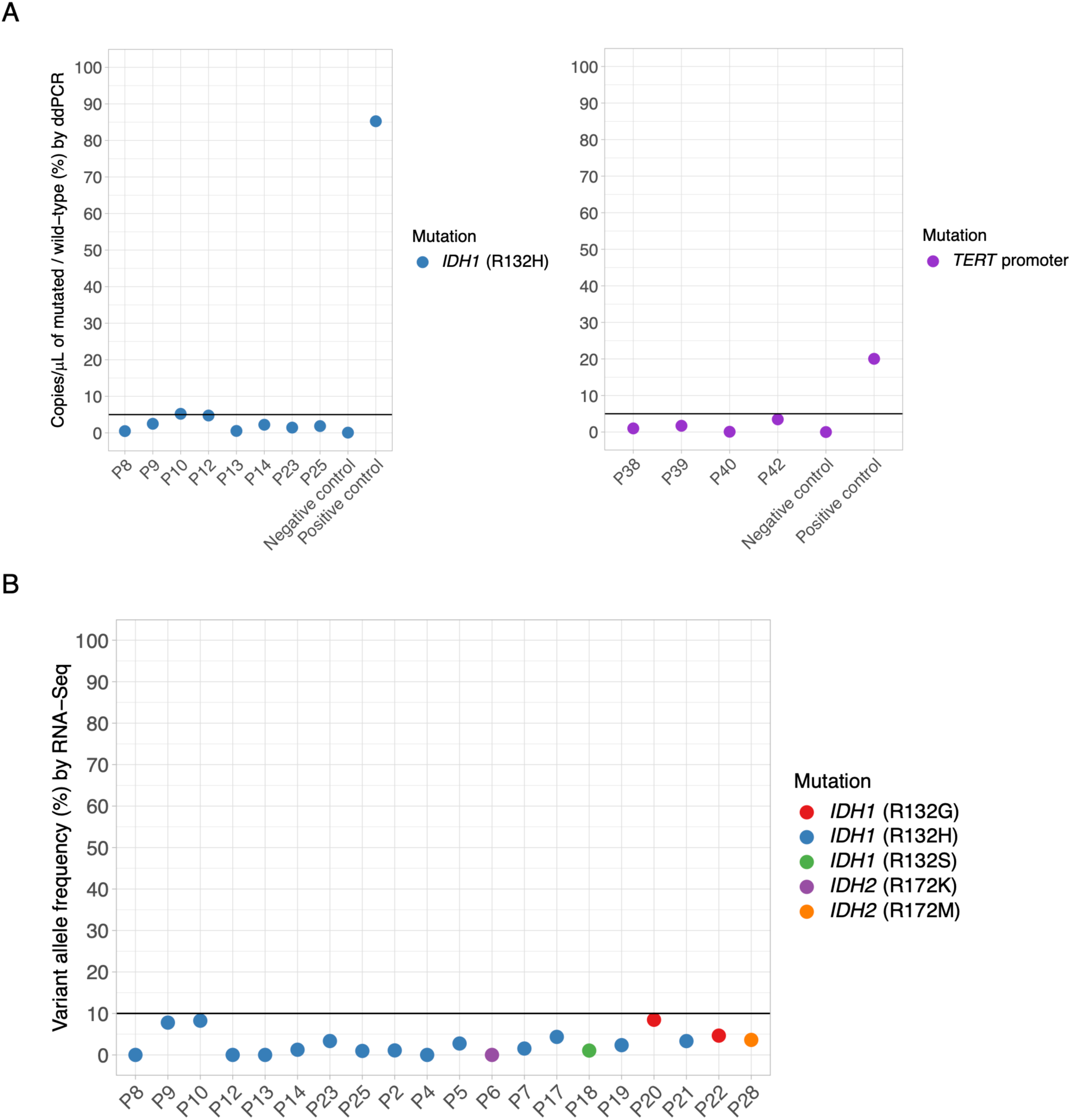
Purity assessment of CD11B^+^ myeloid fractions isolated from human gliomas. **(A)** Digital droplet PCR (ddPCR) using mutation-specific assays for IDH1 R132H and TERT promoter mutations C228T/C250T, together with the corresponding wild-type alleles, performed on available DNA from CD11B+ fractions (*n* = 12) isolated from gliomas with known mutation status. Less than 5% mutant copies/µL relative to the corresponding wild-type alleles were detected in each analyzed sample. Positive and negative bulk-tumor controls for each assay are also shown. Differences in the relative detection of IDH and TERT mutations in positive controls may reflect regional clonal heterogeneity within the tumors. **(B)** Expression frequency of IDH-mutant alleles assessed in RNA-seq data from CD11B^+^ fractions (*n* = 20) isolated from gliomas with a known mutation status using bam-readcount^97^. Compared to ddPCR, RNA-seq-based estimates slightly overestimated tumor-cell contamination in samples P9 and P10

**Supplementary Fig. S2.**
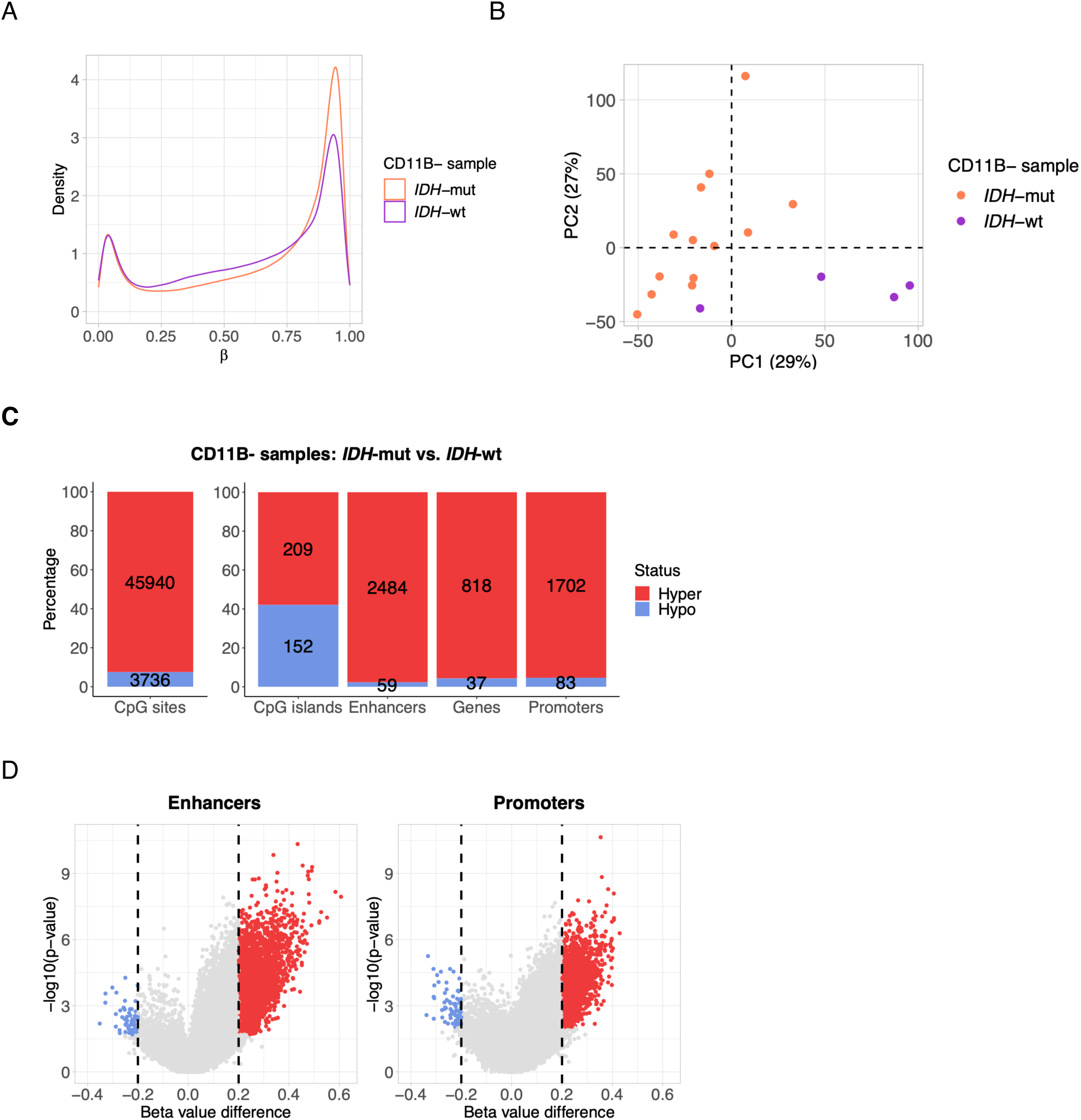
Tumor cells enriched CD11B^−^ fractions display global DNA hypermethylation in IDH-mutant gliomas. **(A)** Density plot showing DNA methylation levels (β-values) in tumor cells enriched CD11B^−^ fractions isolated from IDH-mutant (*n* = 13) and IDH-wt (*n* = 4) gliomas. **(B)** Principal component analysis performed using methylation profiles from approximately 600,000 CpG sites across the genome in CD11B^−^ fractions from the study cohort. **(C)** Stacked bar plots showing the absolute number and relative proportions of differentially methylated CpG sites across four functional genomic regions according to methylation status in CD11B^−^ fractions from IDH-mutant vs. IDH-wt tumors. **(D)** Volcano plots illustrating the magnitude and distribution of differentially methylated enhancers and promoters in CD11B^−^ fractions from the same comparison.

**Supplementary Fig. S3.**
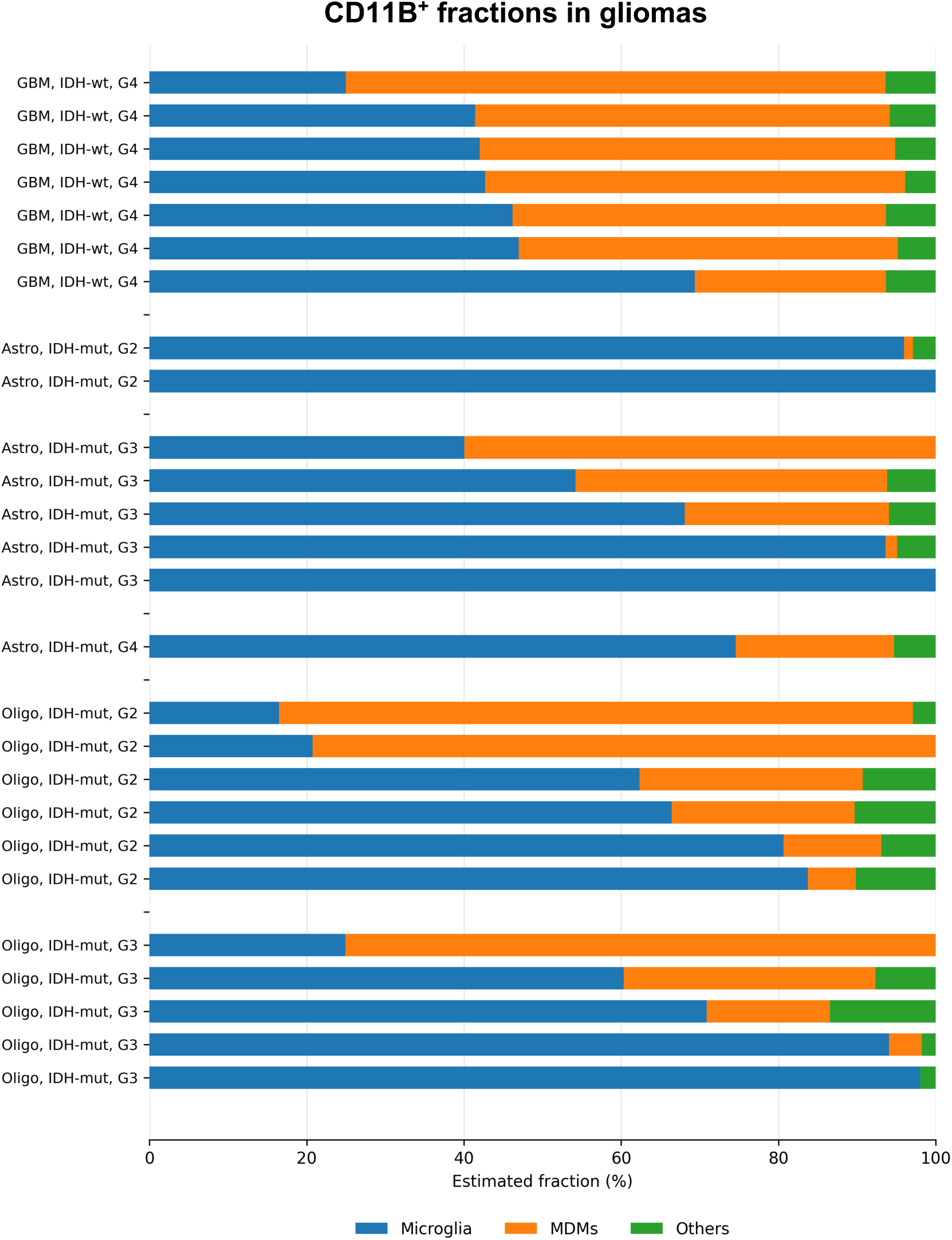
Estimation of microglial and monocyte-derived macrophage (MDM) composition in CD11B⁺ fractions from human gliomas. Methylation-based deconvolution of CD11B⁺ fractions from gliomas, using the Glioma Immune Microenvironment Composition Calculator (GIMiCC)^82^, shows the estimated proportions of resident microglia, monocyte-derived macrophages (MDMs), and other minor myeloid populations (primarily NK cells and granulocytes) in each tumor. Overall, IDH-mutant tumors tended to display higher estimated microglial fractions (∼70%) and lower MDM proportions (∼20%) relative to IDH-wt glioblastomas (∼45–50% microglia and ∼45–50% MDMs). GBM, glioblastoma; Astro, astrocytoma; Oligo, oligodendroglioma; G, CNS WHO grade.

**Supplementary Fig. S4.**
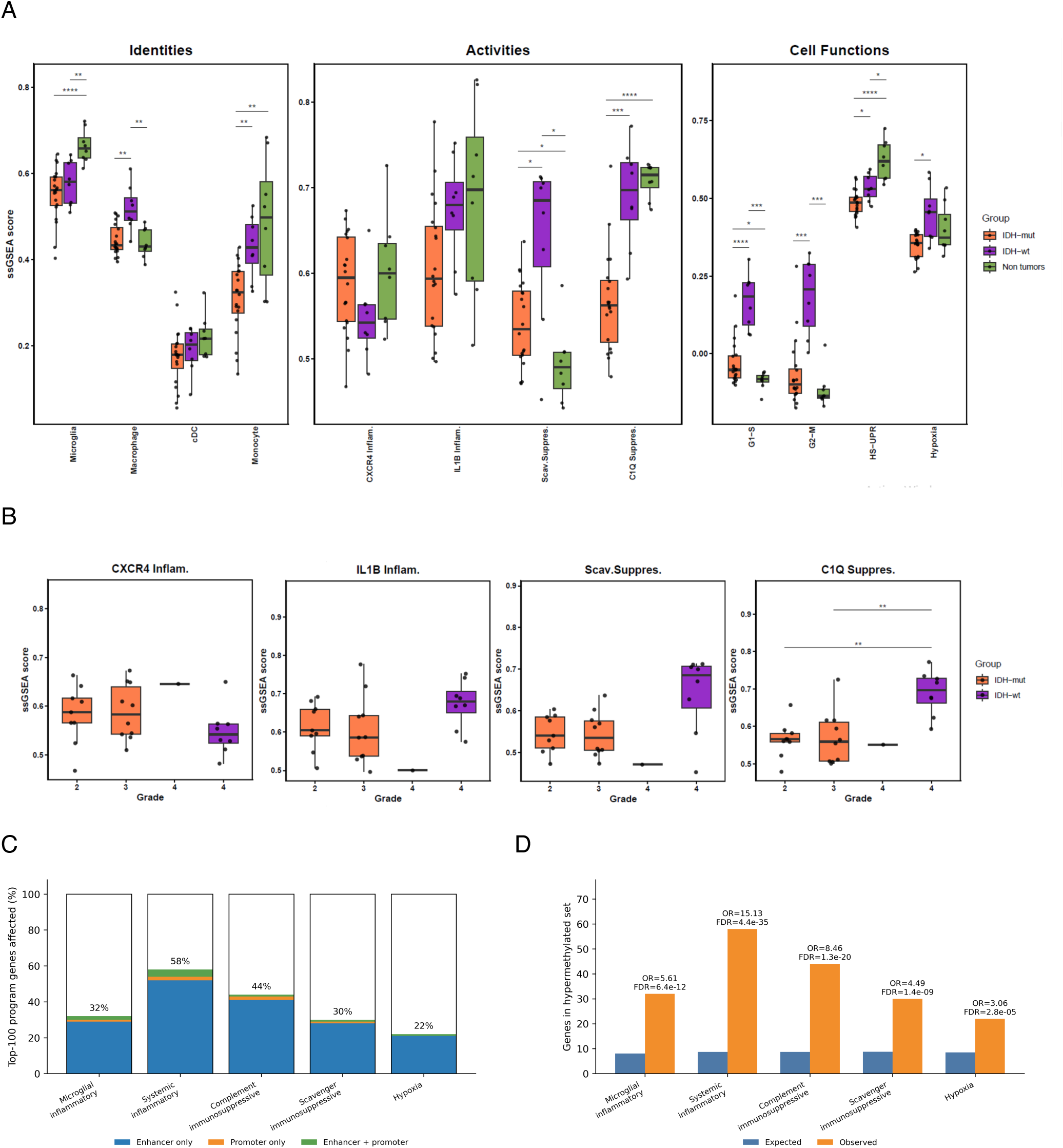
Immunomodulatory programs expressed by CD11B⁺ myeloid cells in gliomas and non-tumor brain tissues. **(A)** Plots showing single-sample gene set enrichment analysis (ssGSEA) scores for the indicated immunomodulatory programs calculated from RNA-seq data. Significance levels correspond to adjusted p-values (*P < 0.05, **P < 0.01, ***P < 0.001; Wilcoxon rank-sum test). **(B)** Same analysis as in (A), stratified to tumor grade. **(C)** Percentage of the top 100 representative genes from each immunomodulatory program associated with hypermethylated regulatory regions in CD11B⁺ fractions from IDH-mutant gliomas. Genes were classified according to whether they were linked to hypermethylated enhancers, promoters, or both. **(D)** Enrichment analysis of genes associated with hypermethylated regulatory regions within the indicated immunomodulatory programs. The tested gene set comprised the union of genes linked to recurrently hypermethylated enhancers and promoters shared across tumor and non-tumor comparisons. One-sided Fisher’s exact tests were performed using the full FANTOM5-based regulatory annotation background employed in the methylome analysis. P-values were adjusted using the Benjamini–Hochberg method. Bars indicate observed and expected numbers of genes within the hypermethylated regulatory region set.

**Supplementary Fig. S5.**
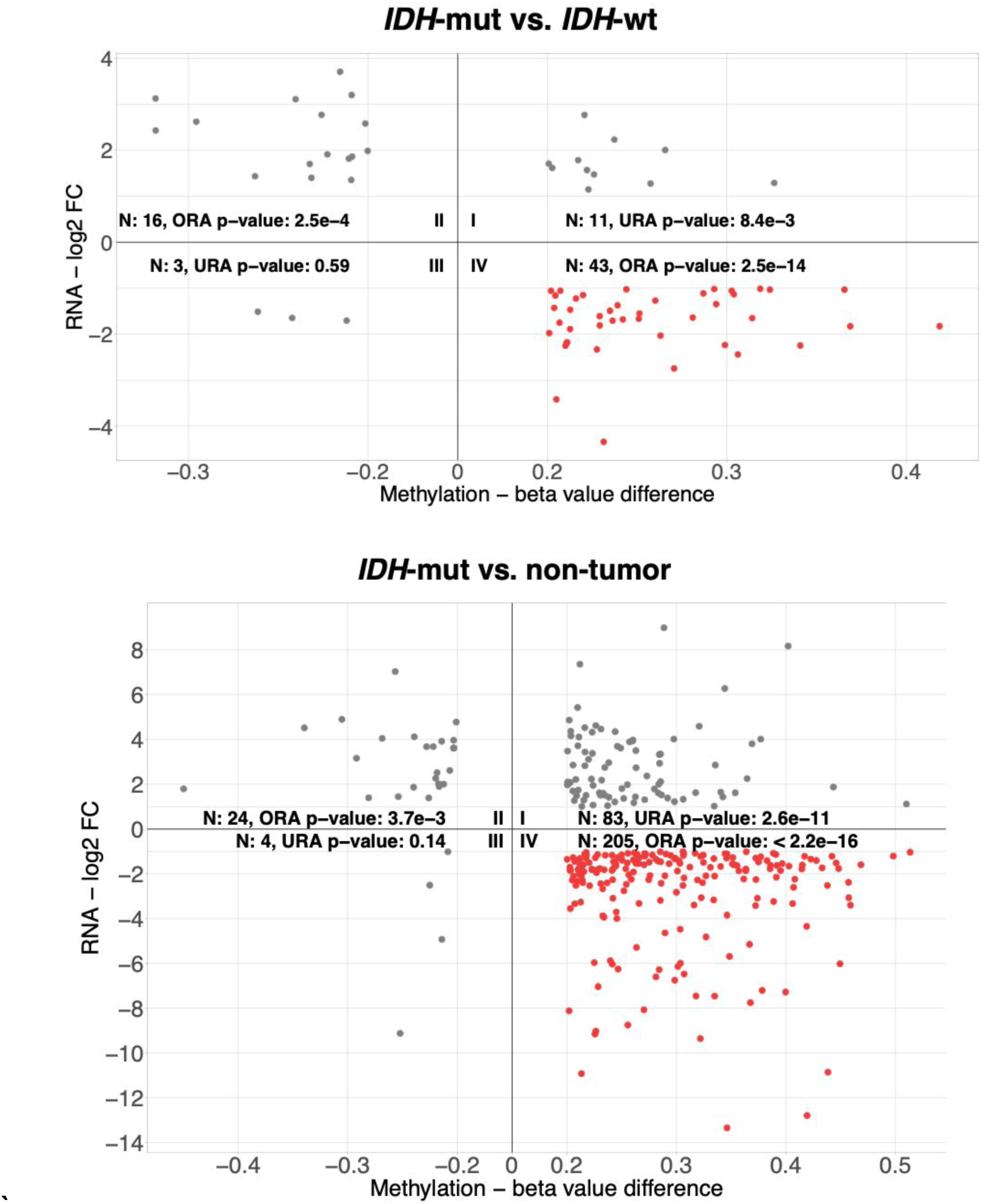
Integrated gene expression and promoter methylation changes in CD11B⁺ myeloid cells from IDH-mutant gliomas across comparisons. Cartesian plots showing genes (dots) exhibiting significant changes in both expression (Y-axis) and promoter methylation (X-axis) in CD11B^+^ fractions from IDH-mutant gliomas compared with IDH-wt tumors (upper panel) and non-tumor brain tissues (lower panel). Numbers indicate the total genes identified in each quadrant. Over-representation and under-representation of canonical methylation/expression relationships (quadrants II and IV) and non-canonical relationships (quadrants I and III) were assessed using over-representation analysis (ORA) and under-representation analysis (URA), respectively. Quadrant IV (highlighted in red) contains genes exhibiting promoter hypermethylation together with transcriptional downregulation.

**Supplementary Fig. S6.**
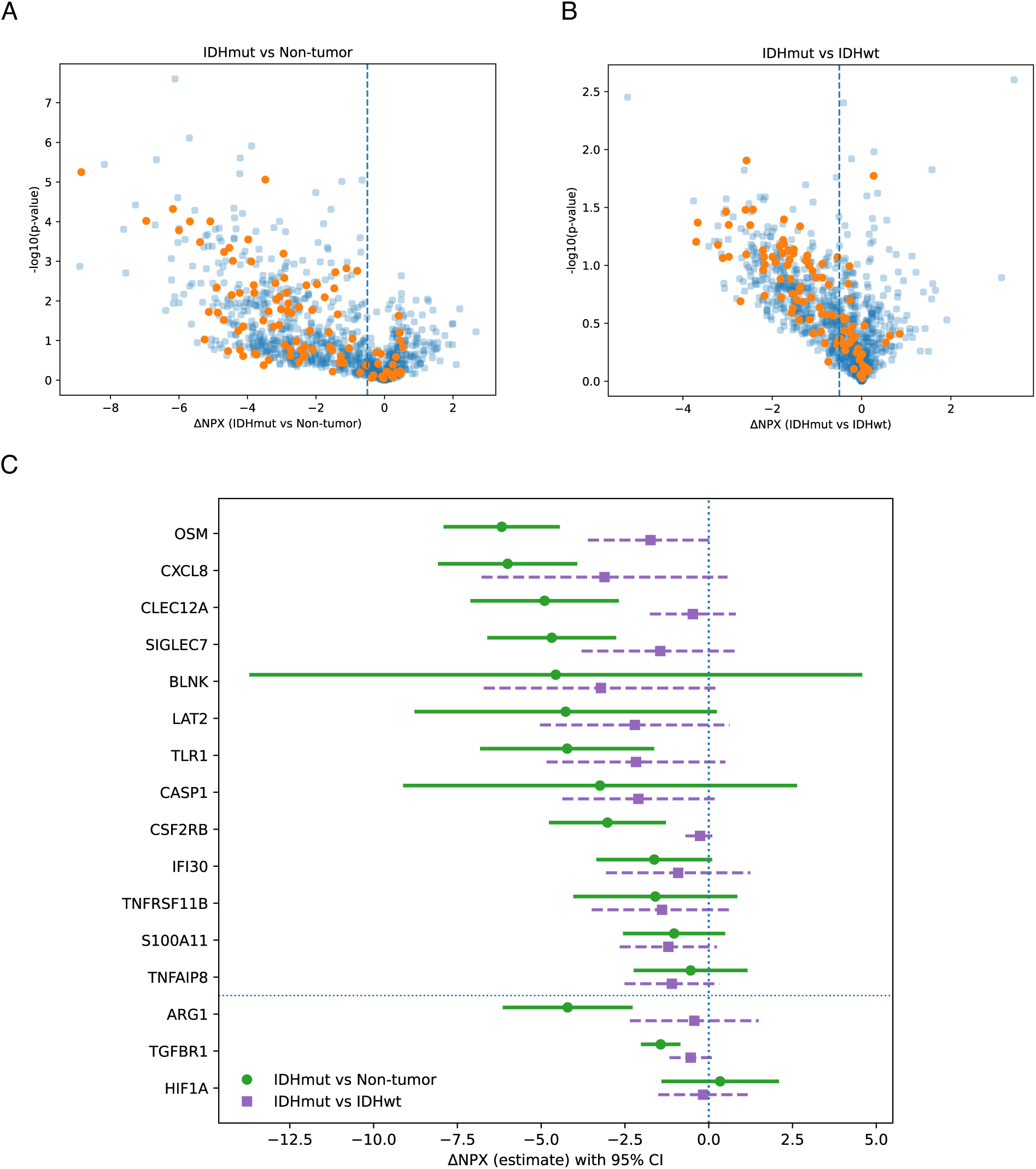
Proteomic validation of enhancer-associated transcriptional repression in myeloid cells from IDH-mutant gliomas. **(A)** Volcano plot showing mean differential protein abundance (ΔNPX) for 1,034 proteins (Olink Reveal panel) in CD11B⁺ fractions from IDH-mutant tumors (*n* = 7) relative to non-tumor brain tissues (*n* = 3). Proteins corresponding to genes identified in the integrative methylation–transcriptome analysis (*n* = 101) are highlighted in orange. The dashed vertical line indicates a biologically meaningful reduction threshold (ΔNPX = −0.5). The 101-protein subset exhibited significantly stronger downregulation than the full panel (median ΔNPX: −2.53 vs −1.19; one-sided Mann– Whitney U test, P = 6.06×10⁻⁶). **(B)** Volcano plot showing the same analysis as in (A) for CD11B⁺ fractions from IDH-mutant (*n* = 7) vs. IDH-wt gliomas (*n* = 6). Changes were directionally consistent but smaller in magnitude. The 101-protein subset remained significantly more downregulated than the full panel (median ΔNPX: −1.05 vs −0.39; one-side Mann– Whitney U test, P = 3.21×10⁻⁶). **(C)** Forest plot showing ΔNPX estimates ± 95% confidence intervals (from Welch’s t-tests) for 13 genes exhibiting concordant enhancer hypermethylation, transcriptional repression, and reduced protein abundance across both comparisons, together with the regulatory comparators HIF1A, ARG1, and TGFBR1. Circles indicate IDH-mutant vs non-tumor comparisons, whereas squares indicate IDH-mutant vs. IDH-wt comparisons. The vertical dotted line indicates no change (ΔNPX = 0). Across the full panel, ΔNPX distributions were significantly shifted below zero in both comparisons (one-sample Wilcoxon signed-rank test against 0; P < 0.001).

**Supplementary Fig. S7.**
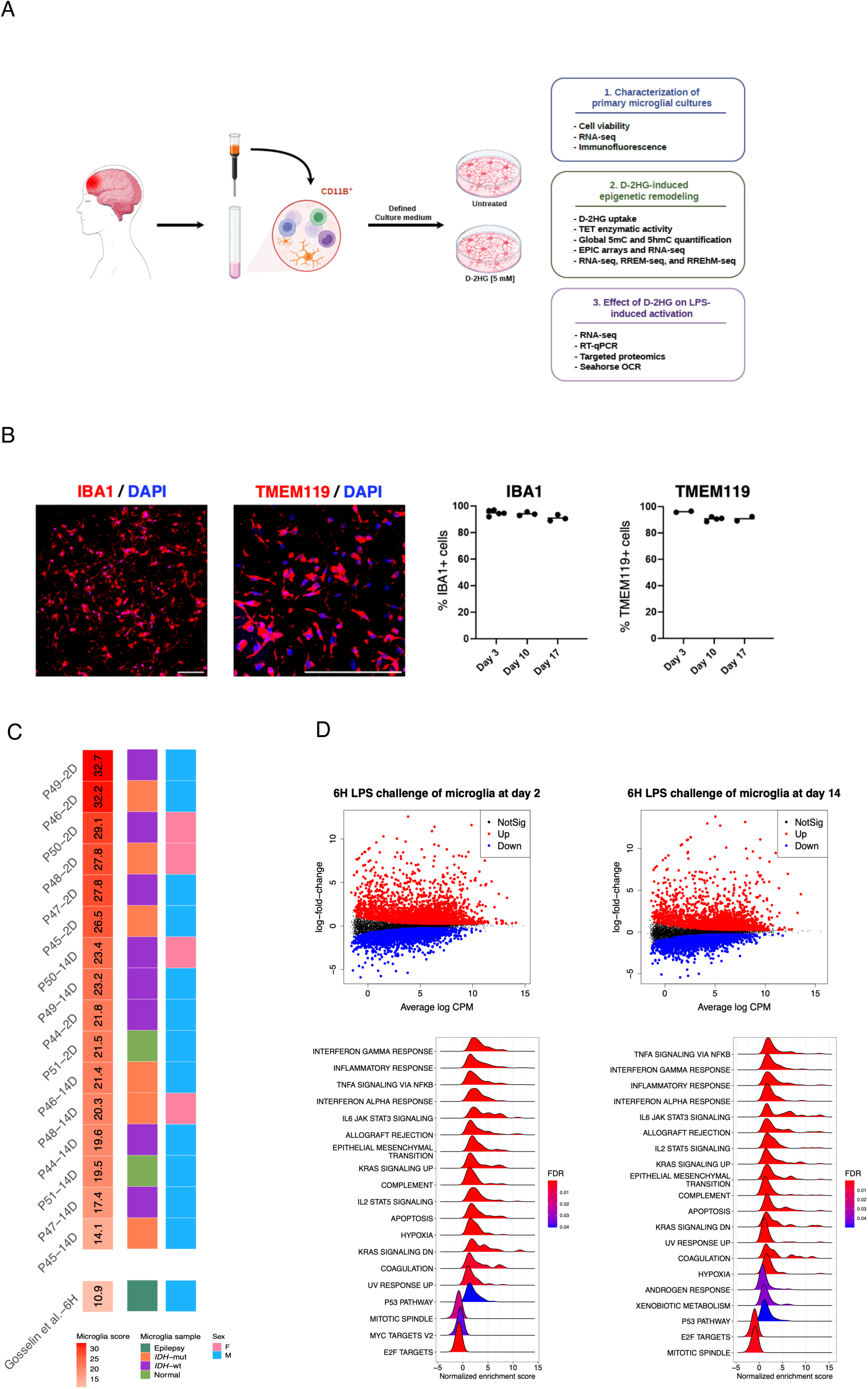
Functional characterization of primary human microglial cultures. **(A)** Overview of the experimental workflow used to establish primary human microglial cultures and the downstream experimental analyses. Tissue collection procedures and culture conditions are described in the Methods section. The assays performed in each primary microglial preparation are indicated in Supplementary Table S1. **(B)** Representative immunofluorescence images showing preserved expression of microglial markers TMEM119 (confocal microscopy) and IBA1 (Apotome microscopy) in primary cultures after 12 days *in vitro* (scale bars: 100 µm). Estimated percentages of TMEM119^+^ and IBA1^+^ cells at different time points are shown on the right. **(C)** Transcriptome assessment (RNA-seq) of microglial identity in primary human microglial cultures (*n* = 8) analyzed after 2 days (2D) or 14 days (14D) *in vitro* using a patient-paired design. Microglial identity scores were calculated using the microglial signature defined by Gosselin et al^39^ from normalized gene-expression matrices. The indicated scores demonstrate preservation of the microglial transcriptional signature even after 14 days in culture under defined conditions, compared with the “culture shock” conditions reported by Gosselin et al. Sample annotations indicating tissue origin and sex are shown below. **(D)** Plots showing differential gene expression in primary human microglial cultures (*n* = 4) analyzed by RNA-seq 6 h after LPS stimulation at 2 days (left) or 14 days (right). Red and blue indicate significantly upregulated and downregulated genes, respectively, (FDR < 0.05), relative to unstimulated controls. **(E)** Ridge plots showing Hallmark gene set enrichment analyses (GSEA) in LPS-stimulated microglia after 2 days (left) or 14 days (right) in culture. Activated (NES > 0) and repressed (NES < 0) gene sets with FDR < 0.05 are shown. **Explanatory note:** Exploratory EPIC-array profiling was performed in a subset of D-2HG-treated primary microglial cultures during the study. These analyses informed the design of subsequent deep-sequencing approaches used to characterize D-2HG–induced enhancer remodeling but are not presented in the current manuscript.

**Supplementary Fig. S8.**
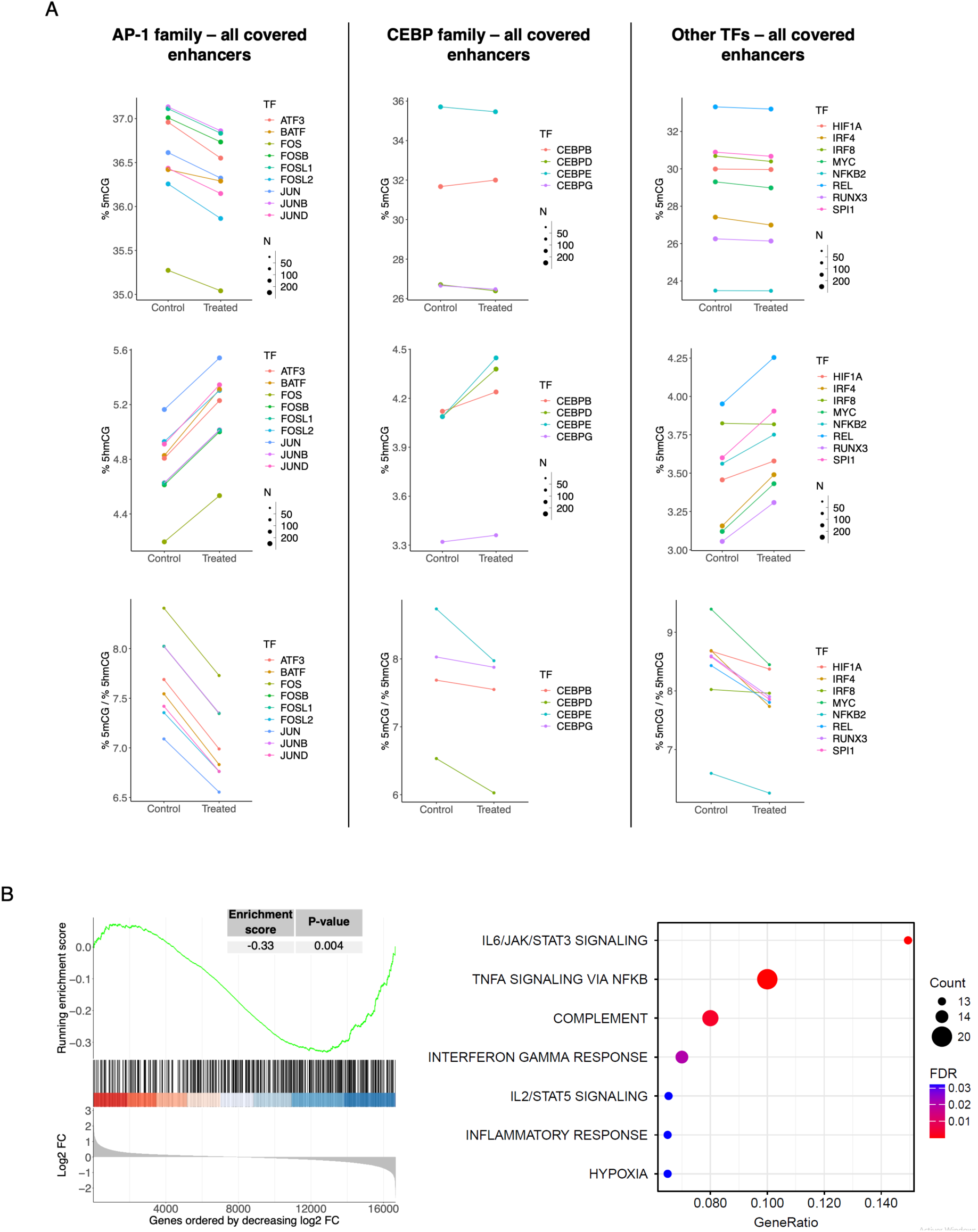
Single-base resolution analysis of 5mC and 5hmC surrounding TFs motifs across all covered enhancers in D-2HG-treated primary microglia. **(A)** Plots showing separate analyses of 5mC and 5hmC percentages (upper panels) surrounding TF-binding motifs located within all covered enhancers for the indicated TFs in D-2HG-treated microglia and paired untreated controls. Dot size indicates the number of motifs analyzed in a paired manner between conditions. Corresponding 5mC/5hmC ratios in the analyzed regions are also shown. **(B)** Gene set enrichment analysis (GSEA) performed using genes ranked by log2FC from D-2HG-treated vs. control RNA-seq samples. The plot shows significant enrichment toward decreased expression of genes associated with D-2HG-driven hypermethylated and hypohydroxymethylated enhancers bound by TFs shown in Fig. 5E. Methylome, hydroxymethylome, and RNA-seq analyses were perform in the same samples using a paired design. **(C)** Bubble plot showing Hallmark gene sets enriched among genes associated with D-2HG-driven hypermethylated and hypohydroxymethylated enhancers. Dot size and color indicate the number of genes contributing to each category and the corresponding FDR value, respectively (FDR < 0.05).

**Supplementary Fig. S9.**
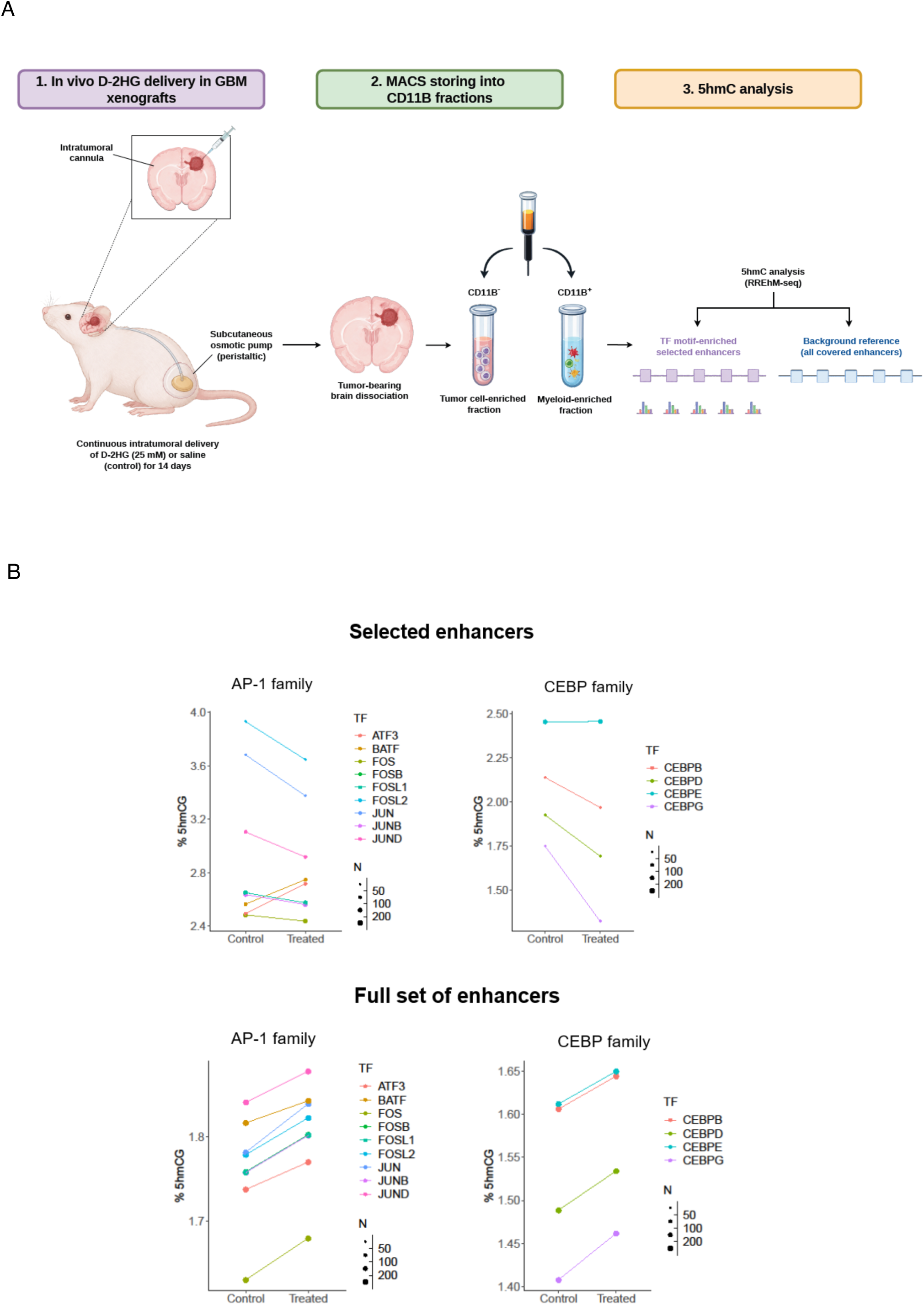
*In vivo* D-2HG delivery model and enhancer hydroxymethylation analyses in CD11B⁺ myeloid cells. **(A)** Schematic overview of the experimental workflow. Nude mice bearing established orthotopic GBM xenografts received continuous intratumoral delivery of D-2HG (25 mM) or saline control for 14 days using subcutaneous osmotic pumps connected to intracerebral cannulas. Tumor-bearing hemispheres were dissociated and CD11B⁺ myeloid cells were isolated by magnetic enrichment prior to RREhM-seq analysis. Due to limited CD11B⁺ cell yields, samples from two mice were pooled prior to sequencing (*n* = 4 mice per condition overall). Single-base resolution hydroxymethylation analyses were performed to interrogate impairment of TET function at enhancer regions. **(B)** Changes in %5hmC at orthologous enhancers enriched for AP-1 and CEBP family TF-binding motifs identified in the *ex vivo* analyses (“selected enhancers”) compared with the broader set of covered enhancers used as a background reference (“full set of enhancers”) in CD11B⁺ myeloid cells isolated from D-2HG-treated and saline-treated mice. Selected enhancers exhibited reduced hydroxymethylation following chronic D-2HG exposure, consistent with the alterations observed in primary human microglia.

**Supplementary Fig. S10.**
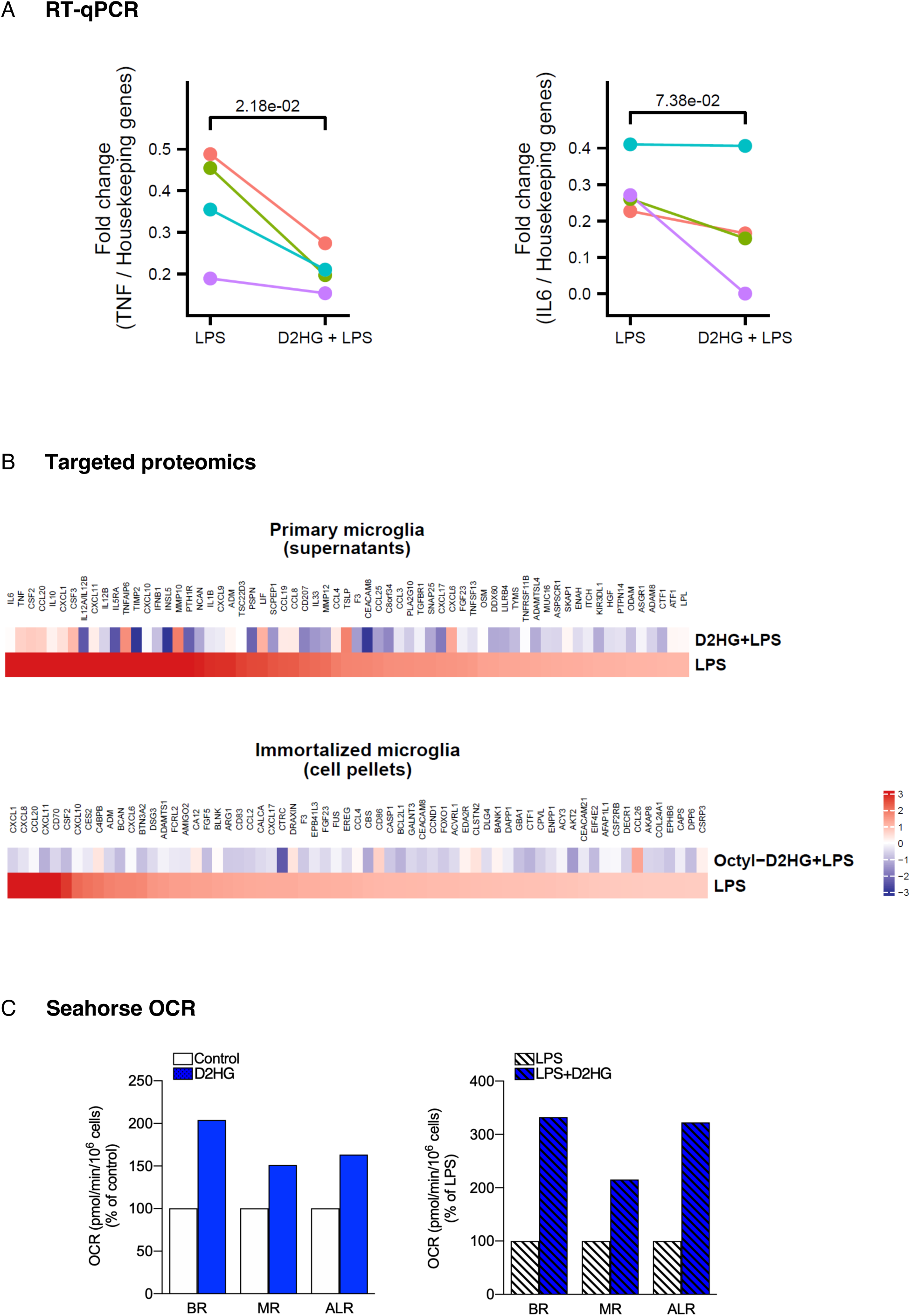
Validation experiments following prolonged D-2HG exposure and subsequent LPS challenge in human microglia. **(A)** Plots showing reduced expression of LPS-induced cytokines in primary microglia (*n* = 4), as determined by RT-qPCR after 14-days of D-2HG treatment. Expression values were normalized to housekeeping genes. P*-*values were calculated using one-sided paired *t*-tests. **(B)** Heatmaps showing reduced induction of LPS-responsive proteins in primary microglia after 14-days of D-2HG treatment or in immortalized microglial cells after 30 days of octyl-D-2HG treatment (*n* = 3), as determined using the Olink Reveal panel in culture supernatants and cell pellets, respectively. **(C)** Assessment of oxygen consumption rate (OCR) in primary microglia using the XFe extracellular flux analyzer, consistent with increased oxidative metabolism following D-2HG exposure. Basal respiration (BR), maximal respiration (MR), and ATP-linked respiration (ALR) were compared between control and D-2HG-treated cells, as well as between LPS- and LPS+D-2HG-treated conditions. All cultures were maintained, treated, and collected under the same conditions as those described for Fig. 6A.

**Supplementary Fig. S11.**
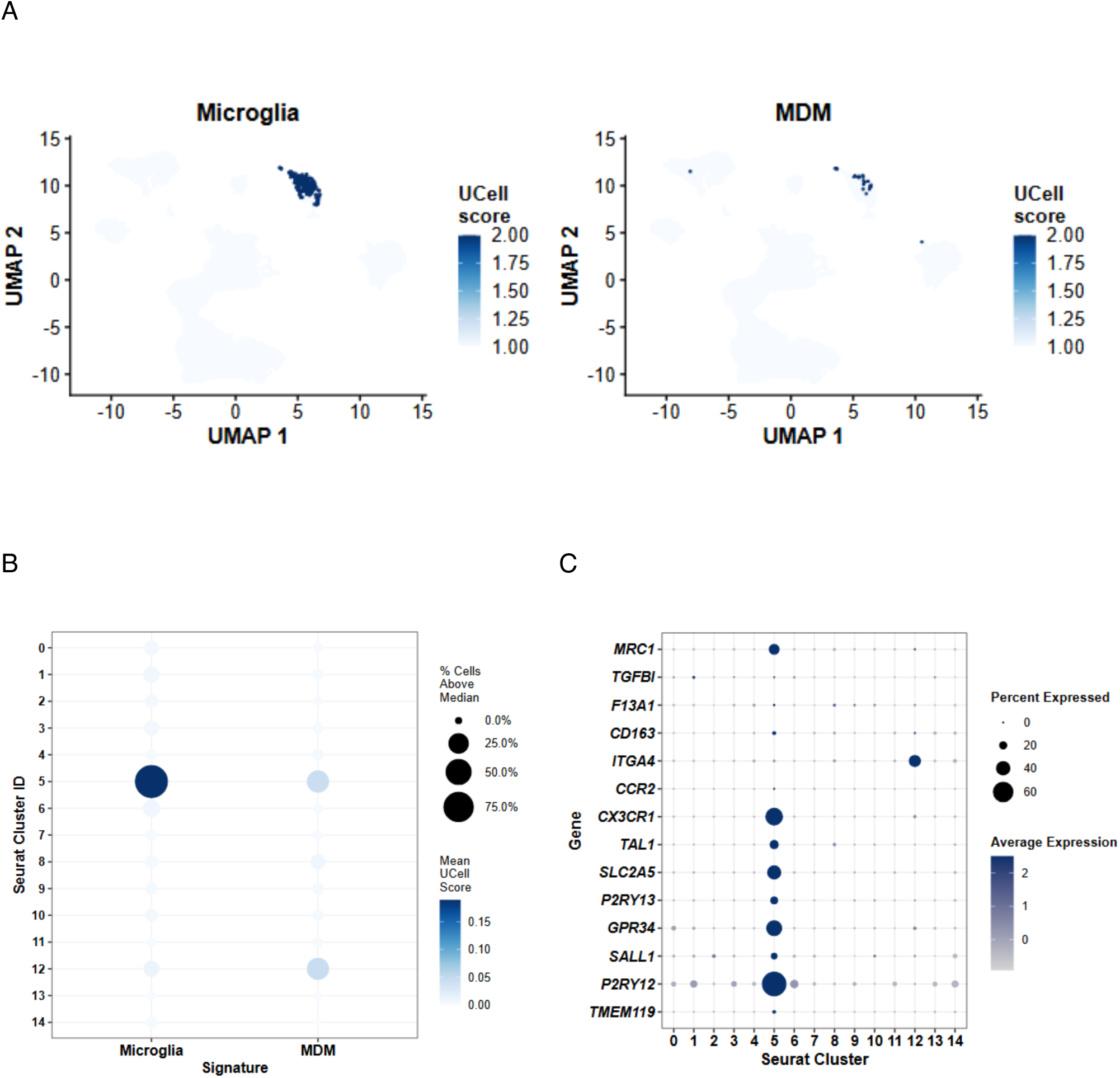
Identification and characterization of microglia in mutant IDH inhibitor–treated patients. **(A)** UMAP representation of the integrated single-cell/nucleus transcriptomic dataset showing UCell scores for curated microglia and monocyte-derived macrophage (MDM) gene signatures. **(B)** Dot plot showing enrichment of microglia and MDM transcriptional signatures across Seurat clusters. Dot size indicates the percentage of cells with UCell scores above the median, and color indicates the mean UCell score. **(C)** Dot plot showing expression of representative microglia (TMEM119, P2RY12, SALL1, GPR34, P2RY13, SLC2A5, TAL1, and CX3CR1) and MDM (CCR2, ITGA4, CD163, F13A1, TGFBI, and MRC1) marker genes across Seurat clusters. Dot size indicates the percentage of expressing cells and color indicates average expression level.

## Supplementary Tables

**Supplementary Table S1.** Clinical, molecular and demographic characteristics of the study cohort and type of assays performed in CD11B^+^ fractions.

**Supplementary Table S2.** Genes associated with hypermethylated enhancers and promoters shared across comparisons in CD11B^+^ fractions from IDH-mutant gliomas.

**Supplementary Table S3.** Differentially expressed genes in CD11B^+^ fractions from IDH-mut gliomas compared with IDH-wt tumors and non-tumor brain tissues.

**Supplementary Table S4.** Hallmark pathway enrichment analyses of upregulated and downregulated genes in CD11B^+^ fractions from IDH-mutant gliomas.

**Supplementary Table S5.** Downregulated genes linked to hypermethylated enhancer regions in CD11B+ myeloid fractions from IDH-mutant gliomas.

**Supplementary Table S6.** Targeted proteomic analyses in CD11B⁺ fractions from IDH-mutant gliomas across comparisons.

**Supplementary Table S7.** Genes associated with enhancers displaying hypermethylation and hypohydroxymethylation in D-2HG-treated primary human microglia.

**Supplementary Table S8.** Gene expression changes in microglia from patients treated with mutant IDH inhibitors assessed by snRNA-seq.

